# Single Cell Transcriptome of Stress Vulnerability Network in mouse Prefrontal Cortex

**DOI:** 10.1101/2023.05.14.540705

**Authors:** Benjamin Hing, Sara B. Mitchell, Maureen Eberle, Yassine Filali, Ian Hultman, Molly Matkovich, Mukundan Kasturirangan, Whitney Wyche, Alli Jimenez, Radha Velamuri, Micah Johnson, Sanvesh Srivastava, Rainbo Hultman

## Abstract

Increased vulnerability to stress is a major risk factor for the manifestation of several mood disorders, including major depressive disorder (MDD). Despite the status of MDD as a significant donor to global disability, the complex integration of genetic and environmental factors that contribute to the behavioral display of such disorders has made a thorough understanding of related etiology elusive. Recent developments suggest that a brain-wide network approach is needed, taking into account the complex interplay of cell types spanning multiple brain regions. Single cell RNA-sequencing technologies can provide transcriptomic profiling at the single-cell level across heterogenous samples. Furthermore, we have previously used local field potential oscillations and machine learning to identify an electrical brain network that is indicative of a predisposed vulnerability state. Thus, this study combined single cell RNA-sequencing (scRNA-Seq) with electrical brain network measures of the stress-vulnerable state, providing a unique opportunity to access the relationship between stress network activity and transcriptomic changes within individual cell types. We found especially high numbers of differentially expressed genes between animals with high and low stress vulnerability brain network activity in astrocytes and glutamatergic neurons but we estimated that vulnerability network activity depends most on GABAergic neurons. High vulnerability network activity included upregulation of microglia and mitochondrial and metabolic pathways, while lower vulnerability involved synaptic regulation. Genes that were differentially regulated with vulnerability network activity significantly overlapped with genes identified as having significant SNPs by human GWAS for depression. Taken together, these data provide the gene expression architecture of a previously uncharacterized stress vulnerability brain state, enabling new understanding and intervention of predisposition to stress susceptibility.

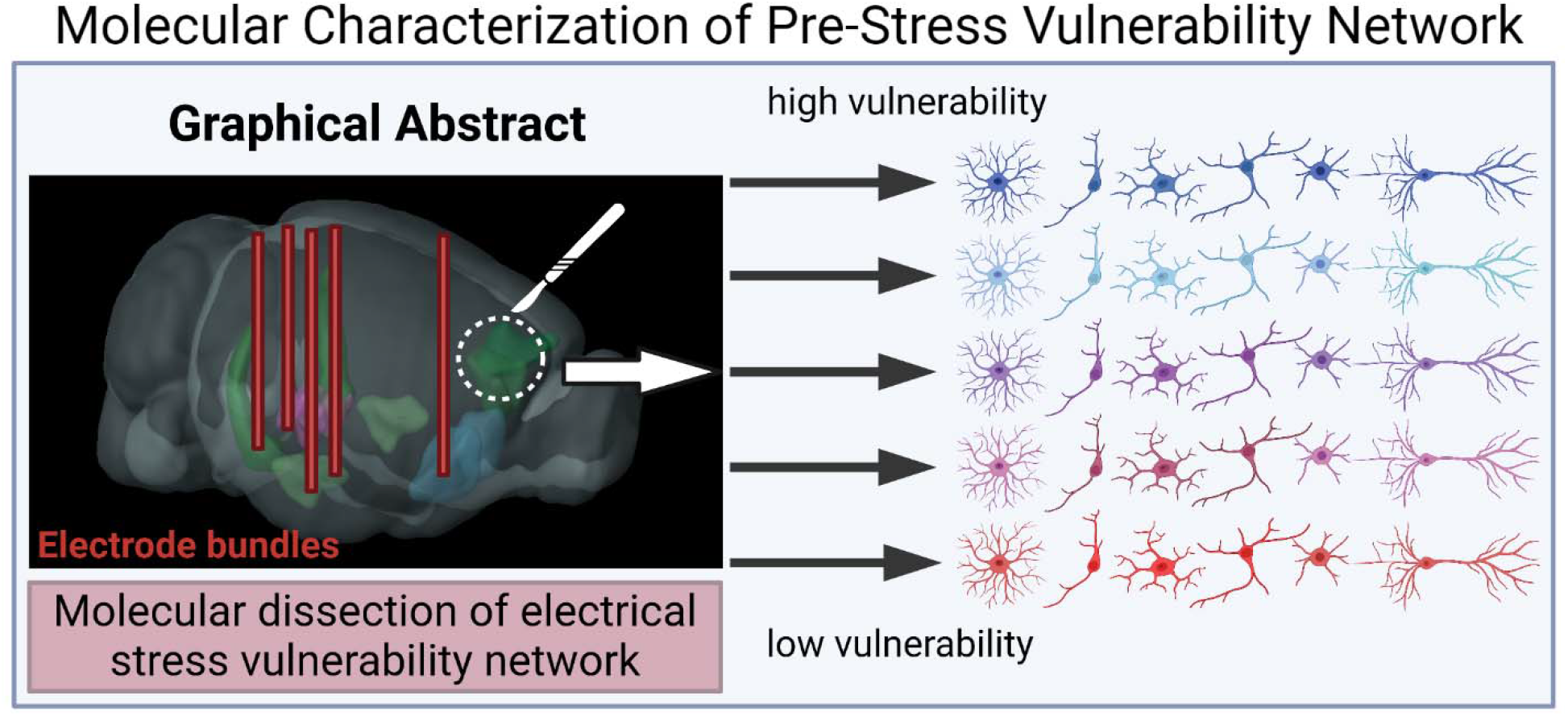

## Intro

Stress has been demonstrated to be a major environmental contributor to myriad disorders, particularly mood disorders such as major depressive disorder (MDD) ^1^. Most such disorders are polygenic, with greater numbers of SNPs increasing an individual’s risk to psychiatric disease. Genes conferring risk work together in complex ways across different cell types, sometimes in near dose-dependent ways, suggesting that a more complete understanding of gene expression events will be necessary for understanding the etiology of such disorders ^2–5^. This appears especially true for MDD, a psychiatric disorder for which heritability is most diverse and susceptible to environmental contributors such as stress ^6,7^.

Toward this end, substantial progress has been made in understanding the gene regulatory events distinguishing stress-susceptible and resilient brain states in rodent models of depressive brain states ^8,9^. In particular, transcriptional regulation of the prefrontal cortex (PFC), has been shown to play an important role in mediating stress susceptibility and in disorders that can often be exacerbated by stress such as MDD ^9–12^. At least five different cell types have been implicated in the etiology of stress susceptibility and related disorders such as MDD in the PFC including excitatory ^13–16^ and inhibitory ^2,17^ neurons, astrocytes ^18^, oligodendrocytes ^19^, and microglia ^20^.

While it has long been recognized that stress has the potential to trigger or exacerbate negative affect states, the gene expression events defining the predisposing vulnerability state has not been rigorously characterized in the same way that post-stress susceptibility and resilience have. Previously, we used multi-site *in vivo* neurophysiology and machine learning methods to identify and validate a local field potential (LFP)-based brain network, *Electome Factor* 1 (*EF*1), that could reliably identify a stress vulnerability state without the full manifestation of a stress-induced depressive state ^21^. Here, we record *EF*1 brain network activity and assess gene expression in specific cell types within the PFC of the same animals, a key driving region of this vulnerability network, to provide a better understanding of the stress-vulnerable state as well as the gene expression architecture of this electrical brain network.

## Results

The overall goal of these experiments was to identify gene expression underlying a previously defined stress-vulnerability brain state, a brain state recognized to predispose animals to manifesting a depressive phenotype following stress ^21^. A neurophysiological metric of the stress vulnerability state was used, *Electome Factor 1*, characterized by local field potential (LFP) features spanning seven brain regions: prelimbic (PrL) and infralimbic (IL) cortices, nucleus accumbens (NAc), basolateral (BLA) and central (CeA) amygdala, ventral hippocampus (Vhipp), and ventral tegmental area (VTA). A complete set of these features is outlined in Figure 1B and Figure S2. Having previously established that the strength of this network signature can be measured over time using a subset of just five brain regions, in the current study we implanted 6 male and 6 female C57BL/6 animals in NAc, BLA, CeA, VTA, and vhipp ^21^. We carried out a forced interaction test (FIT) and calculated *Electome Factor 1* using a previously developed method of projecting new data onto our previously developed stress *Electome Factor* features (Figure 1A, B) ^21^. We then dissected prefrontal cortex (PFC), which included PrL and IL for single cell RNA-Seq (Figure S3).

**Figure 1.**
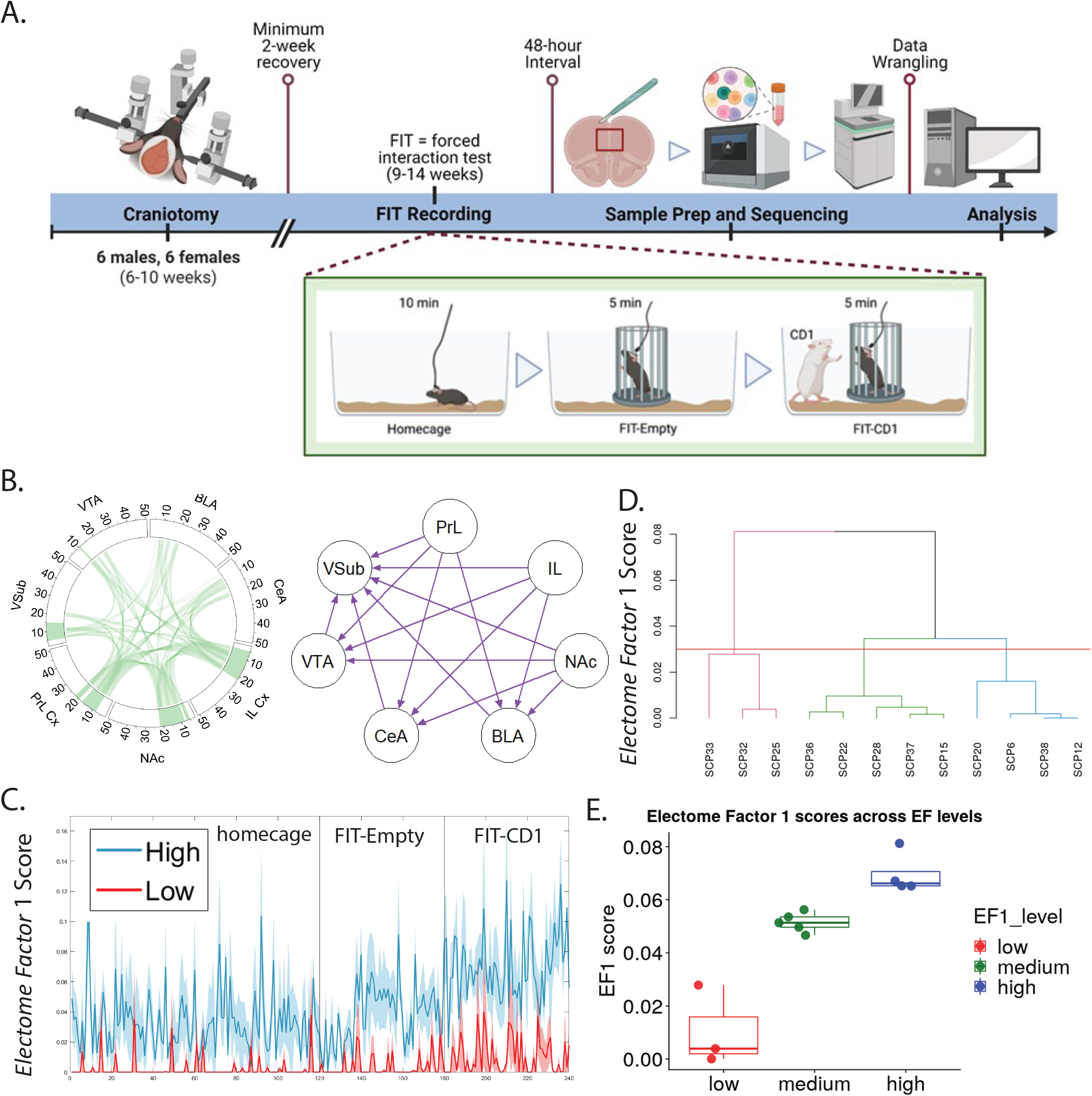
Sorting animals by vulnerability brain states. A) Experimental overview: from left to right implantation of multi-site in vivo electrodes, electrical recordings during forced interaction test (FIT) to measure *Electome Factor* 1 stress vulnerability network, single cell RNA-Seq, bioinformatic data analysis. B) LFP features of *Electome Factor* 1 C) *Electome Factor* 1 activity over time on FIT average of high (blue) and low (pink) clustering animals in bold with SEM in light blue and pink, respectively. D) *Electome Factor* 1 score clusters: low (pink), medium (green), and high (blue).

In order to identify genes that are differentially regulated between animals with high and low stress-vulnerability electrical brain network activity (*Electome Factor* 1), we clustered *Electome Factor* 1 scores using a hierarchical clustering method (Figure 1D). We identified three animals with the lowest *Electome Factor* 1 scores (i.e. corresponding with more stress-resistance) and four animals with the highest *Electome Factor* 1 scores (i.e. corresponding to the more stress-vulnerable, Figure 1E). *Electome Factor* 1 scores over time on the forced interaction test (FIT) are shown in Figure 1C.

### Identification of cell types in the PFC

Single cell analysis identified clusters corresponding to nine major cell types in the PFC (Figure 2A-B, Figure S5). Their distribution was not influenced by factors such as sex, *EF*1 score and experimental dates (Figure S6). The proportion of these cell types were similar between males and females (Figure 2C) and is in line with other studies ^22,23^. While the proportion of these cell types did not differ between high and low *EF*1 groups, GABAergic neurons had a significantly higher proportion (FDR < 0.05) in *EF*1 high group compared to *EF*1 low group (Figure 2D).

**Figure 2.**
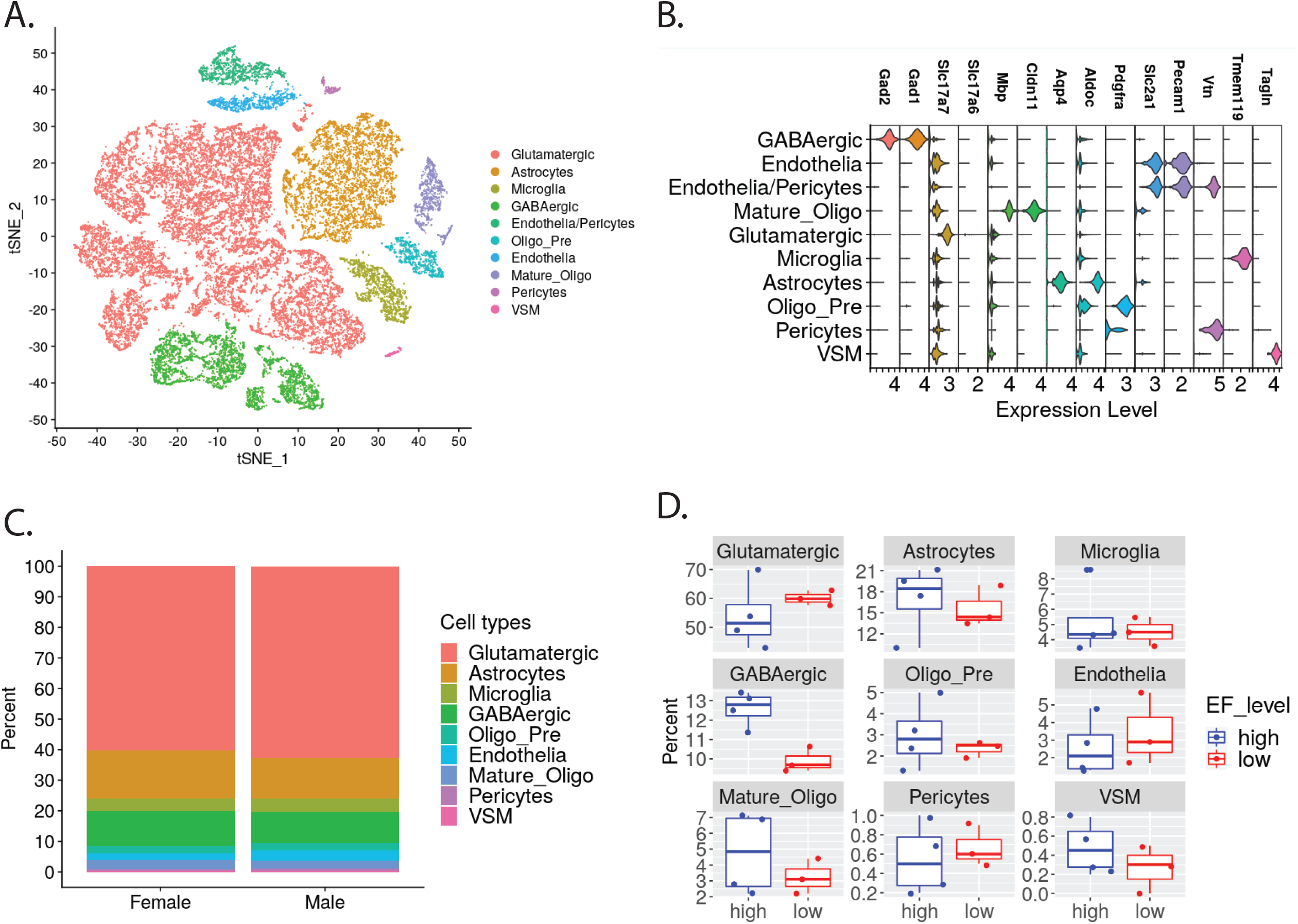
Single cell RNA-Seq cell type distributions. A) t-distributed stochastic neighbor embedding (t-sne) plot of cells isolated from prefrontal cortex (PFC) from electrode-implanted animals. B) violin plot of cell-type specific markers C) cell type distribution by sex D) percentage cell types by high and low *Electome Factor* 1 network activity scores.

### Single Cell Gene Expression Analysis

To evaluate how genes are differentially regulated in the PFC of high and low *EF*1 groups, differential gene expression was performed on cell types which have a minimum of 10 cells for all samples (Table S3). Analysis was performed using the zero-inflated negative binomial model (ZINB)-WaVE-+ DEseq2 workflow. ZINB-WaVE is a flexible scRNA-Seq signal detection method that has been shown to provide accurate and stable representations of data at low dimensions, while eliminating the need for preliminary normalization ^24^. To limit the influence of technical variation in the analysis, mitochondrial derived genes were removed from the data and we used counts corrected for mitochondria percentage obtained from SCTranform for the analysis.

Glutamatergic neurons had the greatest number of significant differentially expressed genes, with 838 genes significantly more highly expressed in the PFC of mice with high *EF*1 scores, (Table S8, Figure 3A, Table S7). Gene ontology (GO) analysis of these genes showed significant over-representation in biological processes including mRNA processing, respiratory processes and mitochondria related processes (Figure 3B and Table S5). In contrast, 548 were significantly more highly expressed in mice with low *EF*1 scores. Analyses by GO showed significant over-representation in biological processes involved with synapse organization and regulation (Figure 3C and Table S5). To further support this result, we performed gene set enrichment analysis (GSEA), which is a threshold-free approach to identifying changes in biological processes between groups ^25^. Consistent with the GO analysis, mice with high *EF*1 scores were significantly enriched for biological processes involved in respiratory and mitochondria related processes (Table S5). Mice with low *EF*1 score did not have significant enrichment for biological processes after correction for multiple testing. The top enrichment terms that trended towards significance included neuronal and synapse related function such as synaptic transmission of glutamatergic (FDR = 0.065), synapse assembly (FDR = 0.066) and glutamatergic receptor signaling pathway (FDR = 0.075).

**Figure 3.**
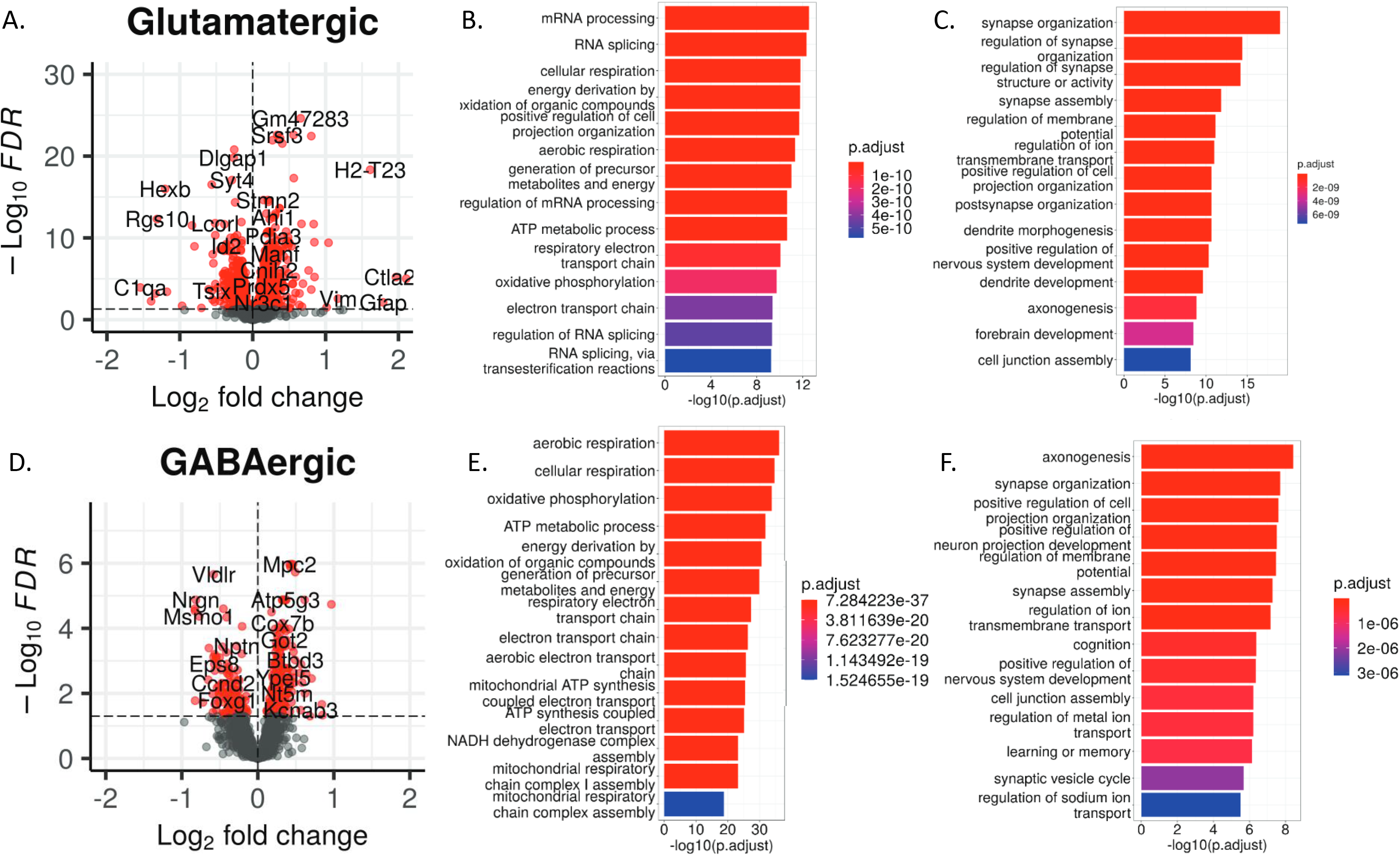
Differential Gene Expression and Gene Ontology Analyses in neurons. Differential gene expression analysis conducted using ZINB-WaVE with sex as a covariate: A) volcano plot of differential gene expression between high and low *Electome Factor* 1 (stress vulnerability) scores in glutamatergic neurons B) gene ontology (GO) analysis for genes upregulated with high/low *Electome Factor* 1 scores in glutamatergic neurons C) gene ontology (GO) analysis for genes downregulated with high/low *Electome Factor* 1 scores in glutamatergic neurons D) volcano plot of differential gene expression between high and low Electome Factor 1 (stress vulnerability) scores in GABAergic neurons E) gene ontology (GO) analysis for genes upregulated with high/low *Electome Factor* 1 scores in GABAergic neurons F) gene ontology (GO) analysis for genes downregulated with high/low *Electome Factor* 1 scores in GABAergic neurons.

The cell type with the second-highest number of DEGs was astrocytes. There were 341 genes significantly highly expressed in mice with high *EF*1 (Table S8, Figure 4A). GO analysis showed significant over-representation of these genes in biological processes including ribosomal and mitochondria related processes (Figure 4B and Table S5). Conversely, there were 221 genes significantly highly expressed in mice with low *EF*1 scores. Analysis by GO showed over-representation of these genes in neuronal related processes including various synaptic processes, regulation of synapse, neurotransmitter secretion and transport (Figure 4C, Table S5). In line with the GO analysis, GSEA also showed significant enrichment of respiratory, and mitochondria related processes in the high *EF*1 group (Table S5). GSEA also supported the findings of the GO analysis in the low *EF*1 group, as there was significant enrichment of neuronal related processes including synapse organization, assembly and synapse signaling (Table S5).

**Figure 4.**
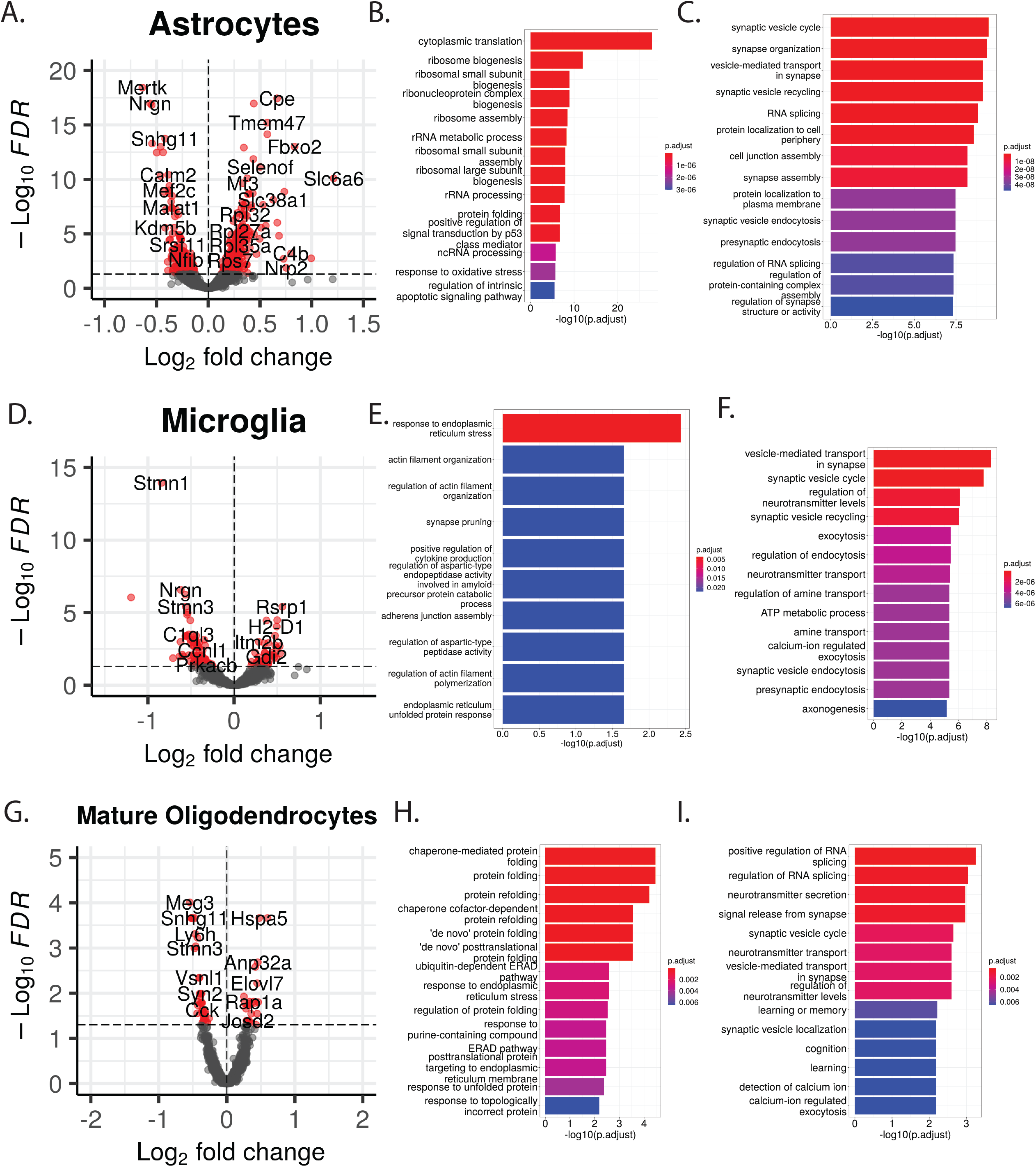
Differential Gene Expression and Gene Ontology Analyses in glial cells. Differential gene expression analysis conducted using Zinbwave with sex as a covariate: A) volcano plot of differential gene expression between high and low *Electome Factor* 1 (stress vulnerability) scores in astrocytes B) astrocyte GO paths upregulated with higher *EF*1 scores C) astrocyte GO paths upregulated with lower *EF*1 scores D) volcano plot of differential gene expression between high and low *Electome Factor* 1 (stress vulnerability) scores in microglia E) microglia GO paths upregulated with higher *EF*1 scores F) microglia GO paths upregulated with lower *EF*1 scores G) volcano plot of differential gene expression between high and low *Electome Factor* 1 (stress vulnerability) scores in mature oligodendrocytes H) mature oligodendrocyte GO paths upregulated with higher *EF*1 scores I) mature oligodendrocyte GO paths upregulated with lower *EF*1 scores

Differential gene expression analysis of GABAergic neurons identified 242 genes that were significantly more highly expressed in the high *EF*1 mice (Table S8, Figure 3D). As with analyses of glutamatergic neurons and astrocytes, analysis of genes significantly more highly expressed with *EF*1 in GABAergic neurons by GO revealed significantly over-representation of these genes in biological processes involved in cellular respiration, oxidation, metabolism and mitochondria related processes (Figure 3E, Table S5). By comparison, there were 140 genes significantly highly expressed in the low *EF*1 group. Biological processes evaluated by GO analysis showed significant over-representation of these genes in neuronal related processes including axonogenesis, synapse organization and assembly, ion transport and cognition (Figure 3F, Table S5). This result was also verified by GSEA which showed significant enrichment of terms for the high *EF*1 group which includes cellular respiration and mitochondria related processes (Table S5). Similarly, the low *EF*1 group showed significant enrichment for neuronal related processes such as regulation of synapse function, synapse assembly and synapse organization (Table S5).

Differential gene expression was performed on microglia, which showed 46 genes significantly more highly expressed in the high *EF*1 group (Table S8, Figure 4D). While GO showed over-representation of these genes in processes involved in the endoplasmic reticulum stress response and synapse pruning, it also highlighted processes involved in proinflammation such as cellular response to IL7, lymphocyte mediated immunity and regulation of actin filament organization (Figure 4E, Table S5). In contrast, there were 77 genes with significantly higher expression in the low *EF*1 group. Analysis by GO showed a significant over-representation of these genes in processes involved in the regulation of synapse activity including synapse vesical cycle, synapse vesical cycle, synaptic vesical endocytosis and regulation of neurotransmitter levels (Figure 4F, Table S5). In addition to GO analysis, GSEA also observed significant enrichment of cellular respiration and mitochondria related function in the high *EF*1 group (Table S5). By comparison, the low *EF*1 group had significant enrichment of processes involved in neuronal processes such as the generation of neurons, developmental growth, and neurogenesis (Table S5).

Finally, we also evaluated differential gene expression of mature oligodendrocytes. We observed 23 genes that were significantly highly expressed in the high *EF*1 group (Table S8, Figure 4G). Analysis by GO showed over-representation in processes involved in protein folding (Figure 4H, Table S5). By comparison, there were 37 genes significantly highly expressed in the low *EF*1 group (Figure 4I, Table S5) which were over-represented in processes involved in neuronal transmission processes including neurotransmitter secretion, signal release from synapse and regulation of neurotransmitter levels. GSEA analysis showed significant enrichment in processes involved in cellular respiration, metabolic processes and mitochondrial function for the high *EF*1 group (Table S5). In contrast, there was an enrichment of neuronal processes including cell signaling, neuron generation and development in the low EF1 group (Table S5).

Given that mitochondria related function was enriched in the high *EF*1 group across all cell types, we sought to further scrutinize this result by performing GSEA using a curated mitochondria dataset from MitoCarta which is a database that catalogs mitochondria related proteins and pathways ^26^. We reasoned that if our findings of mitochondrial significance were truly specific to the high *EF*1 state, we would only see mitochondrial enrichment in pathways for the high *EF*1 group and not for the low *EF*1 group. GSEA showed enrichment of mitochondria related pathways in all five cell types mentioned above in the high *EF*1 group (Table S6). There was however no enrichment in the low *EF*1 group.

### Validation of differential gene expression analysis

To verify that our approach of using ZINB-WaVE + DEseq2 analysis to identify differentially expressed genes is sound, we performed differential gene expression analysis using an independent program DESingle (Table S12, Figure S9), an R package that utilizes a zero-inflated negative binomial model to distinguish between actual zero expression and technical dropout zeros for the detection of various DEGs in scRNA-Seq data with improved accuracy ^27^. Not only did the DEGS detected by DESingle align with ZINB-WaVE + DEseq2 in terms of the ranked quantity of DEGs detected by cell-type (Table S10), but the DEGs identified from the ZINB-WaVE + DEseq2 also largely overlapped with those detected by DESingle analysis (Figure S10). Using DEGs detected by DESingle, GO analysis of genes more highly expressed in the high *EF*1 group showed similar over-representation of biological processes as reflected in our initial analysis including metabolism and mitochondria related processes (Figure S11, Table S13) for glutamatergic neurons, GABAergic neurons, and astrocytes. By comparison, GO analysis of genes more highly expressed in the low *EF*1 group showed over-representation in processes involved in synaptic functions including synapse organization, assembly, axonogenesis, and synaptic vesicle cycle for all the cell types (Figure S11, Table S13).

When GO analysis was performed using genes that overlapped between both analysis methods, we observed a similar result whereby genes more highly expressed in the high *EF*1 group were over- represented in processes such as cellular respiration and mitochondria processes for glutamatergic neurons, GABAergic neurons and astrocytes (Table S14). There was also an over-representation of these genes in ribosomal processes in astrocytes. For microglia, there was over-representation of genes involved in proinflammation and immunoactivation processes including positive regulation of tumor necrosis factor, antigen processing and presentation of peptide antigen, positive regulation of interleukin 6 and positive regulation of phagocytosis (Table S14). In contrast, genes more highly expressed in the low *EF*1 group were over-represented for processes involved in regulation of neuronal or synaptic activity in all major cell types (Table S14).

Given our analysis had shown enrichment for processes involved in synapse regulation in the low *EF*1 group across all cell types, we sought to exclude the possibility of a platform-dependent finding by validating this further by performing an over-representation analysis using SynGO - a database containing expert curation of genes/proteins pertaining to the ontologies for the synapse ^28^. Consistent with our GO and GSEA analysis, we observed significant over-representation of genes highly expressed in the low *EF*1 group for biological processes and cellular components related to synapse function for all cell types (Figure S12 F-J). In contrast, this over-representation was observed to a lesser degree or not observed for genes more highly expressed in the high *EF*1 group for the different cell types (Figure S12 A-E), providing strong corroboration that expression of genes related to synaptic function are indeed upregulated in the low *EF*1 state.

### Upstream Regulator Analysis

Given the differences in gene expression between *EF*1 groups for all the cell types, we further explored upstream regulators that might be involved in the transcriptional differences using Ingenuity Pathway Analysis (IPA) (Table S15). For each transcription factor any available predicted activation states are listed. For glutamatergic neurons, IPA predicted many upstream regulators to be associated with observed DEGs. The top 10 most significant transcriptional regulators out of 70 identified are HTT, *TP53, NFE2L2* (activation*), MYC, TEAD1* (activation), *MYCN, KDM5A* (inhibition), *RB1* (activation), *LARP1*, and *CREB1* (activation). Similarly for GABAergic neurons, the top 10 upstream transcriptional regulators out of 62 identified include *TEAD1* (activation), *KDM5A* (inhibition), *HTT, PPARGC1A* (activation), *NRF1* (activation), *RB1* (activation), *MYC* (activation), *TP53, CREB1* (activation), and *SMYD1*.

For non-neuronal cell such as astrocytes, the top 10 upstream transcriptional regulators out of 80 identified were found to include MLXIPL (activation), *CTNNB1, MYCN* (activation), *MYC* (activation), *YAP1, HTT, TP53, RRP1B*, and *FUS*. Similarly for microglia, the top 10 transcriptional regulators out of 162 identified were predicted to be *HTT, TP53, MYC, Fus, MYCN, NFE2L2, PAX3, HDAC4, PIAS1*, and *NFKBIA*. For mature oligodendrocytes, the top 10 upstream transcriptional regulators out of 38 identified were *HTT, FOXO4* (inhibited), *NFE2L2*, Rest (activated), *SREBF1, CREB1, MYCN, CALR*, *TP53,* and *Fus*.

Overall, while several predicted top upstream transcriptional regulators were consistently present across the different cell types, some transcriptional regulators were present for only particular cell types. For example, *HTT* and *TP53* appear within the top 10 most significant transcriptional regulators for all five cell types being evaluated, while the transcriptional regulators *LARP1, PPARGC1A, MLXIPL*, *PAX3*, and *FOXO4* were exclusively linked with the top 10 most significant transcriptional regulators for Glutamatergic neurons, GABAergic neurons, astrocytes, microglia, and mature oligodendrocytes respectively.

### Cell-Type–*EF*1 Relationships

In an effort to understand which cell types might be contributing most to *EF*1 score activity, we used a generalized additive model (GAM) approach (see Supplemental Results, Figures S13-S18). Due to the large number of gene expression values compared to the number of mice, we used GAM to estimate the non-linear relationships between the principal components of cell-specific gene expressions and the *EF*1 activity. We found that the *EF*1 activity is strongly related to the PCs specific to the GABAergic cells, with the astrocytes-specific PCs also demonstrating some relationship with *EF*1 activity (Table 1, Figure S19). As such, we focused several of our more detailed analyses on these two cell types.

**Table 1.** Cell-type–EF1 relationships as determined by generalized additive model (GAM) Approximate p-value and adjusted R-squared for each GAM model’s estimate of the relationship between fourth-root transformed *EF*1 scores and a single combination of cell type with principal component.

| GAM Modeling Results |  |  |  |
| --- | --- | --- | --- |
| cell label | princ comp | p_val | R2 |
| Astrocytes | PC1 | 0.037090711 | 0.762787933 |
| Astrocytes | PC2 | 0.436914377 | 0.048762311 |
| GABAergic | PC1 | 0.021816254 | 0.859856942 |
| GABAergic | PC2 | 0.019327998 | 0.977880559 |
| Glutamatergic | PC1 | 0.372881787 | 0.132494891 |
| Glutamatergic | PC2 | 0.069872873 | 0.303857349 |
| Mature_Oligo | PC1 | 0.730770506 | 0 |
| Mature_Oligo | PC2 | 0.268270173 | 0.037742471 |
| Microglia | PC1 | 0.52646961 | 0 |
| Microglia | PC2 | 0.421790507 | 0 |

### Comparison of differentially expressed genes with those from chronic stress paradigm

Since vulnerability increases the risk for a depressive phenotype (i.e. “susceptibility”) following chronic stress, we sought to compare our data to post-stress data from the PFC. Our rationale was that identifying genes that are more highly expressed in a pre-stress vulnerability brain state may help to identify which genes are already differentially regulated prior to stress and thus more likely to be causal than compensatory to the effects of stress. We compared our DEGs to those obtained from mice susceptible or resilient to chronic social defeat stress (CSDS) in Bagot, R. et al ^9^. This study evaluated post-stress transcriptional differences in gene expression between susceptible and resilient animals at three time points: 48 hours after chronic stress (early), 28 days after chronic stress (late), and 28 days plus one hour after an additional bout of stress (primed). It also identified larger co-expression networks of susceptibility and resilience. We observed few to no DEGs that overlap with our dataset for susceptible mice that were assessed 48 hours after CSDS, nor with mice primed with previous CSDS exposure and that underwent an additional session of CSDS. However, at 28 days after CSDS, we observed 11 genes for glutamatergic neurons, 6 for GABAergic neurons, 31 for astrocytes and 2 genes for mature oligodendrocytes that overlapped with our dataset (Table S7, Summary_Overlap_with_Bagot, column C, cell-type specific DEG gene lists, columns J. Comparing our DEGs against co-expression networks of resilience and susceptibility, from across all three revealed substantial overlap across many of the modules for both susceptible and resilient networks (Tables S18 and S19. For example, the most striking overlap, with 880 shared genes, was observed between our Glutamatergic neuron DEGs and the genes present in the turquoise resilience module, which is enriched in mitochondrial genes (Table S19, Glutamatergic Sheet, Column B). Furthermore, when evaluated for key driving “hub” genes in this study, several overlapping genes were identified between the Bagot et al., midnightblue susceptible-specific hub genes and our DEGs by cell type, offering possible genes that might regulate an entire network prior to stress (Table S20).

### Comparison of Differentially Expressed Genes (DEGs) with human Genome-Wide Association Studies (GWAS)

In order to assess the potential translational relevance of our findings, we compared genes we found to be significantly differentially expressed with stress vulnerable brain network activity against human gene variants associated with brain disorders. We hypothesized that there would be a significant overlap between genes associated with major depressive disorder and our DEGs related to stress vulnerability, as CSDS has been most closely described to disease as a depression model. Knowing that many psychiatric disorders are polygenic and related, we also evaluated the relationship between our DEGs and similar psychiatric disorders. As a control for specificity, we also evaluated overlap between our DEGs and an unrelated disorder, Alzheimer’s Disease. We reasoned that the nature of the relationships between variants and human disease is variable, with some contributing an increase and others a decrease in gene function and therefore it would be of interest to see genes either up- or down- regulated with vulnerability scores. Most notably, we found that genes with differentially lowered expression with *EF*1 network activity was strongly associated with MDD, bipolar disorder, and schizophrenia and not significantly associated with any other disorders (Figure 5, Table S21). All five cell types evaluated demonstrated some overlap between DEGs and psychiatric disorders (MDD, bipolar, SCZ) but no significant overlap was detected for ADHD, Alzheimer’s, Anxiety/panic disorder, OCD, or PTSD (Figure 5). These results suggest some biological specificity between genes involved in the regulation of stress vulnerability network activity and stress-sensitive depressive mood disorders.

**Figure 5.**
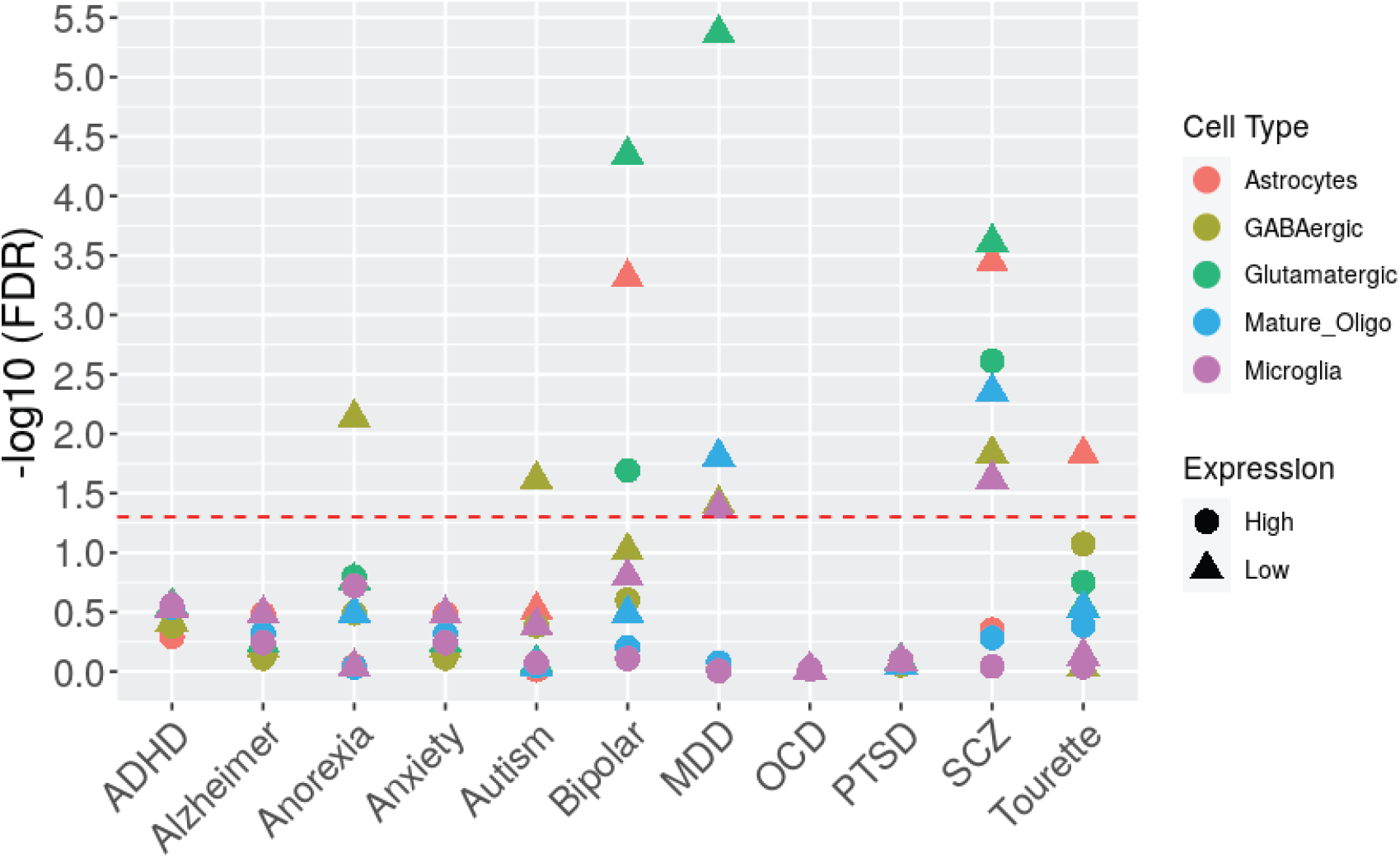
MAGMA analysis to compare differentially expressed genes according to EF1 activity in mice with GWAS datasets from different brain disorders. Genes that were significantly more highly expressed in mice with high vulnerability scores (1; High) or significantly reduced in mice with high vulnerability scores (*▲*; Low) from different cell types were compared with GWAS data from different PGC datasets. Red dotted line represents FDR = 0.05.

### Identification of gene co-expression networks

Given the broad transcriptional changes between *EF*1 high and low groups for different cell types, we sought to determine how networks of genes are organized to contribute to such brain states by performing high dimensional Weighed Gene Correlation Network Analysis (hdWGCNA). This method is particularly useful since small changes in co-regulated genes can contribute to phenotypic states ^29^. Initially, we integrated gene expression data across all cell types and constructed two independent networks, one for *EF*1 high and one for low *EF*1 low; these networks showed broad differences between *EF*1 groups (Figure 6A and 6B), reflecting different gene expression networks engaged in the maintenance of each activity pattern. Having identified that the underlying gene expression architecture is different between animals with high and low *EF*1 activity, we next set out to identify one network across which the same modules could be compared for both high and low *EF*1 animals by performing network construction integrating both datasets (Figure 6C). Fifteen modules were identified with their eigengene expression present in various cell types (Figure S21). The eigengene is a representative expression profile of a module defined by the first principal component of the module ^30^. While the eigengene of the black and magenta modules had positive expression in all cell types, modules such as pink had positive expression in GABAergic and glutamatergic neurons (Figure S20). Other modules such as the brown module had the highest positive eigengene expression in GABAergic neurons (Figure S20). Notably, the level of eigengene expression is not necessarily indicative of the impact of that network on *EF*1 activity by that particular cell type. Some of these modules were enriched with DEGs more highly expressed in the high *EF*1 group (Figure 6D). For GABAergic neurons, the pink module was the most highly enriched for DEGs followed by the brown module (Figure 6D, Figure S22A). Similarly for glutamatergic neurons, the pink module had the highest enrichment of DEGs followed by the black and magenta modules (Figure 6D, Figure S22B). For astrocytes, although the pink module was enriched for DEGs, the greenyellow module had the highest enrichment for these genes (Figure 6D, Figure S22C). Similarly for microglia, while the greenyellow module was enriched for DEGs, the green module was the most enriched for DEGs (Figure 6D, Figure S22D).

**Figure 6.**
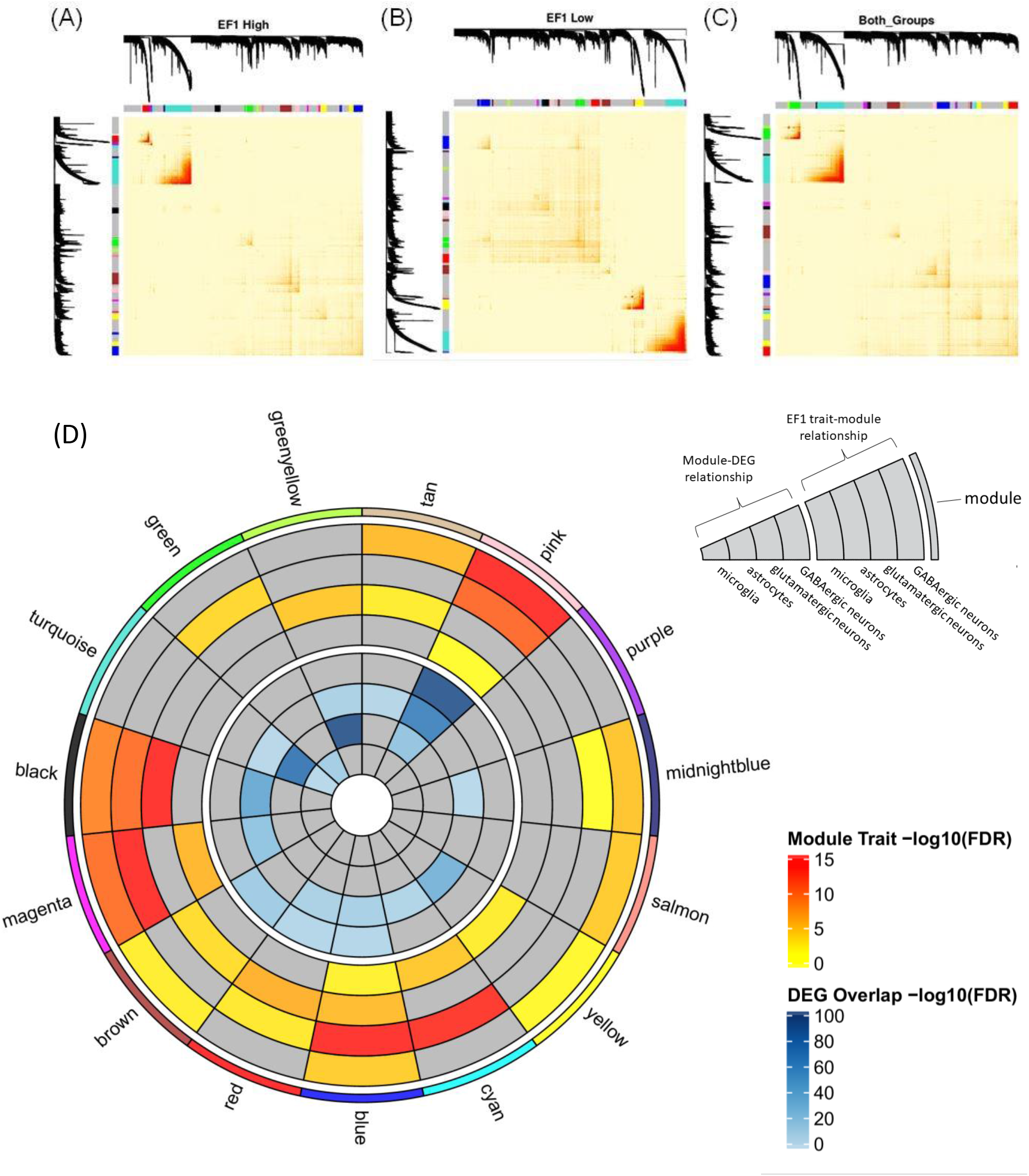
WGCNA analysis to identify gene expression networks of vulnerability network activity. Multi cell type (astrocytes, GABAergic, glutamatergic and microglia) co-expression network was generated for (A) high *EF*1, (B) low *EF*1 and (C) Combined *EF*1 groups. Each panel consists of a topological overlap matrix plot with increasing color intensity of dark red indicating greater topological overlap. On the left and top of each top are dendrograms based on average linkage hierarchical clustering of genes followed by modules below that with arbitrary color assignment colors for identification. (D) Circle plot indicating *EF*1 relationships to different module networks per cell type. Outermost circle indicates each color module, next inner rings indicate module-trait relationships using *EF*1 as the trait across each cell type, innermost rings indicate overlap between module genes and *EF*1 DEGs. Wedge indicate which ring is which cell types.

### Correlation between modules and high and low *EF*1 groups

Since modules are clusters of highly interconnected genes that can regulate phenotypes, we performed module-trait correlation to determine which module is significantly correlated to *EF*1 groups. Notably, eigengene expression from the blue module was significantly correlated with *EF*1 for all cell types (Figure 6D, Figure S20). In contrast, eigengene expression of the pink and magenta modules were significantly correlated with *EF*1 for both GABAergic and glutamatergic neurons and microglia (Figure 6D, Figure S20) while eigengene expression from the black module was significantly correlated with *EF*1 for both neuronal types and astrocytes (Fig. 6D/ Figure S20). For the brown module, its eigengene expression was significantly correlated with *EF*1 for astrocytes and GABAergic neurons for the brown modules (Fig. 6D/ Figure S20).

### Differential eigengene expression (DEE) analysis by *EF*1

Given that the pink module has the highest overlap with DEGs for glutamatergic and GABAergic neurons, we further assessed if this would alter the overall expression of this module as reflected by changes in its eigengene expression between *EF*1 groups. The eigengene of the pink module was significantly more highly expressed in the high *EF*1 group for both neuronal cell types consistent with their DEG expression (Figure S24A). Gene ontology analysis showed that the pink module was enriched for genes involved in biological processes involving mitochondria function and cellular respiration (Figure 7A, Table S22). The functional annotation of other modules is described in Table S22 and Figure S23. No modules were observed to be differentially expressed for microglia (Figure S24D).

**Fig. 7.**
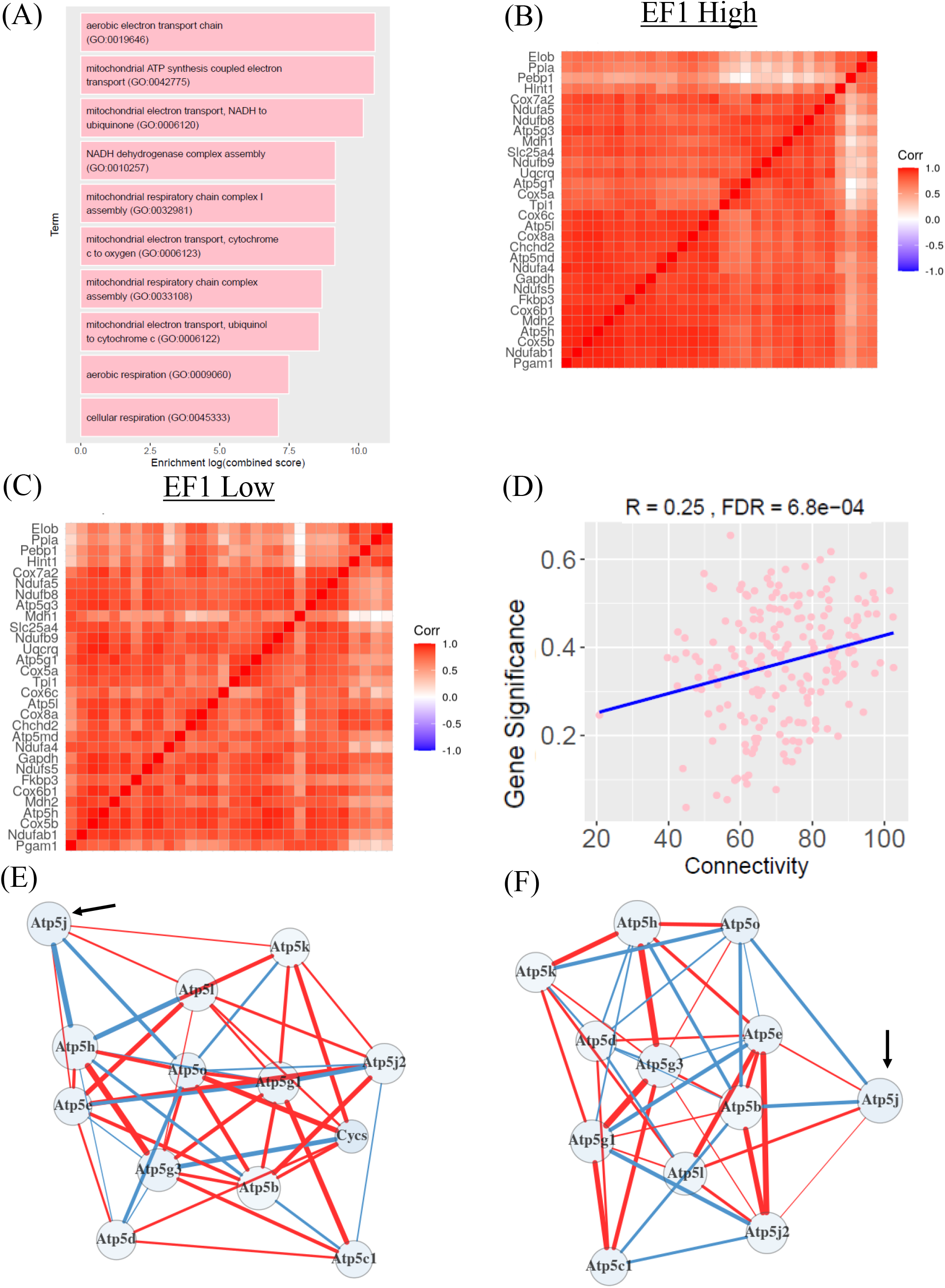
Gene ontology and differential network analysis of the pink module from GABAergic neurons. (A) Gene ontology analysis of biological processes for the pink module. The top 10 terms arranged by combined score that survived multiple correction are shown. (B-C) Network structure was compared between (B) high *EF*1 group and (C) low *EF*1 group using the top 30 genes of the pink module ranked by eigengene-based connectivity from GABAergic neurons. (D) Pearson correlation between intramodular connectivity and Gene significance (absolute value of the correlation between gene and trait) for the pink module. (E) The top gene in the pink module showed differential connectivity for the mitochondrial biogenesis pathway and (F) ATP by chemiosmotic coupling pathway in the pink module. Key gene is highlight with a black arrow for (E,F) Atp5j. Blue nodes indicate higher gene expression in *EF*1 high group. Size of node indicate expression change. Blue line indicates stronger association with high *EF*1 group and red line indicates stronger association with low *EF*1 group. The thickness of the line indicates the strength of association. Figure only show genes in pathways with differential connectivity. FDR; false discovery rate.

### Evaluating differences in network structure between high and low *EF*1 groups

Given that *EF*1 activity is strongly related to PCs specific to GABAergic neurons (Table 1), we evaluated differences in its network structure between *EF*1 groups using discomod ^31^. We focused on the pink module as it is highly relevant to *EF*1 activity. It not only has positive eigenegene expression in GABAergic neurons, it is also associated with *EF*1 group in GABAergic neurons and it module eigengene is differentially expressed between *EF*1 high and low group. Using the top 30 genes ranked by eigengene-based connectivity, we observed significant differences (FDR < 0.002) in correlation value for the pink module between *EF*1 high (Figure 7B) and *EF*1 low (Figure 7C) for GABAergic neurons (Figure 7). To evaluate how changes in network structure alters connectivity in pathways regulated by these modules with regard to *EF*1 activity, we used dnapath ^32^ to assess for changes in connectivity in gene regulatory pathways for this module and observed significant differential connectivity in GABAergic neurons for pathways involving mitochondria function such as mitochondrial biogenesis and formation of ATP by chemiosmotic coupling (Table S23). Differences in connectivity for other modules that associated with *EF*1 groups for GABAergic neurons are described in (Table S23).

### Identification of key genes in pink module

To identify highly relevant modules and genes involved in *EF*1, we reasoned that these modules would have genes with expression that are significantly correlated with *EF*1 (i.e. Gene Significance). Gene significance was previously defined as the absolute correlation value between gene expression and trait^30^. In addition, genes that are important drivers of these networks would also have a high degree of connectivity to other genes. Using this criterion, we observed that the pink module from GABAergic neurons have a significant positive linear relationship between Gene Significance and intramodular connectivity (R = 0.25 respectively, FDR < 0.001) (Figure 7D).

Given that the eigengene expression of the pink module is positively expressed in GABAergic neurons and it shows the greatest relevance for *EF*1, we further identified genes in this network which have the highest connectivity since previous studies have shown that genes that have high connectivity can be influential over the activity of the network ^9^. We identified the Atp5j whose intramodular connectivity was in the top 10^th^ percentile for intramodular connectivity and top 15^th^ percentile for Gene Significance in GABAergic neurons in the pink module for GABAergic neurons. This gene was observed to have differential connectivity between *EF*1 groups in pathways such as mitochondrial biogenesis and formation of ATP by chemiosmotic coupling (Figure 7A). Only one other gene, *Gapdh*, was also observed to have a similar level of intramodular connectivity and Gene Significance and was differentially connected between *EF*1 groups.

## Discussion

Here we have implemented a novel approach combining multi-site in vivo neurophysiology with single cell transcriptomics to provide what to our knowledge is the first single cell resolution molecular characterization of a stress vulnerability brain state. We focused our study on the PFC both because of our previous findings of the importance of the PFC to the electrical network (*EF*1) defining the latent vulnerability state and because of the important role that the PFC plays in regulating stress responses, which suggested to us that understanding its gene regulation in an early stress-vulnerable state may be particularly illuminating ^21,33^. Notably, the differential gene expression identified here does not represent a response to a stressor or molecular manipulation but rather endogenously pre-existing differences between brains with more or less stress vulnerability electrical brain network activity. Nonetheless, we identified robust gene expression differences in these endogenous high and low *EF*1 activity states across all five of the major cell types we examined. Furthermore, we found significant overlap between our differentially expressed genes and human GWAS genes for major depressive disorder (MDD) across all cell types, providing strong evidence of the biological and potential translational relevance of our findings. These overlapping genes also provide a starting point to refine focus amongst the myriad of risk genes involved with MDD for evaluating potential contributors to stress vulnerability.

Beyond the effects observed across all cell types, several observations pointed toward a particular contribution of GABAergic cells to the stress vulnerability brain state defined by *EF*1. For one, GABAergic cells were the only population of cells with significantly different percentage representation between high and low *EF*1 groups (Figure 2D). When considering the overall contribution of different cell types to *EF*1 scores, we used a General Additive Model with principal component values for each cell type. While most cell types demonstrated no obvious relationship with *EF*1 this way, both principal components 1 and 2 of GABAergic cells demonstrated a notably stronger, statistically significant relationship with *EF*1, with astrocytes appearing to have some relationship as well (Supplemental Methods and Results, Table 1 and Figure S19). This finding suggests that GABAergic neurons of the PFC may be the cell type with the most robust contribution to a given animal’s *EF*1 score, and thus, predictive of their pre-stress vulnerability. Given the important role that inhibitory interneurons have been demonstrated to have both in regulation of LFPs, and in stress response and susceptibility, these results were not surprising. Due to the number of cells capture in this study, we did not have sufficient number of sub-types of GABAergic neurons to further explore their contribution to *EF*1 activity. Future studies would require higher number of target cells for such an analysis.

Many rodent studies have shown that early life experiences and adverse exposures can impact GABAergic processes in the PFC, including maturation ^34^, inhibitory circuit development with the BLA and associated perineuronal net formation ^35^, and the programming of GABA transmission ^36^. Along these lines, it has been found that chronic stress affects both the structure and function of prefrontal GABAergic networks ^37,38^, with certain GABAergic sub-types showing enhanced vulnerability to stress ^39^. Interestingly, and in further support of this point, we found trending differences in observed percentage of two GABAergic neuron sub-types: Pvalb and Vip, according to high and low *EF*1 score (Figure S8).

Furthermore, there is myriad additional evidence supporting GABAergic neuronal features playing an important role in mediating stress resilience and susceptibility such as expression level of specific GABA receptors ^40^, signaling peptides ^41^, and subtypes ^17^, as well as interactions with the stress response pathway ^42 43^ and the gut-vagus-brain pathway ^44^. Although the aim of this study was to evaluate PFC cell-type specific gene expression in relation to a brain-wide network signature of pre-stress latent vulnerability, the broad association of GABAergic neurons in the PFC with stress and behavior provides considerable support for GABAergic neurons being a key contributor to *EF*1 activity.

Generally, however, the overall data suggests that it is the coordination of gene expression coming from multiple cell type network components and not one in particular that are responsible for generation of the *EF*1 network. This is supported from our DEG analysis that different cell types had differences in gene expression between *EF*1 groups as well as network analysis. Network analysis showed that modules with expression that spanning different cell types are correlated with *EF*1 groups. Given the important role of multi-brain region networks in both transcriptional and neural activity regulation of stress states ^9,21^, future studies will be aimed at further mapping gene expression by electrical network across multiple brain region states.

In line with this idea of coordinated multicellular network contributions to the vulnerability brain state, our network analysis helped examine the possibility that biological functions represented by modules can extend across cell types and that certain modules may play a bigger role in *EF*1 network. For instance, out of all the modules, the pink module was most enriched for DEGs across three of the four cell types investigated (i.e. GABAergic and glutamatergic neurons, and astrocytes) and genes in this module are enriched for mitochondria related processes which is relevant to the function of different cell types. Its eigengene was also differentially expressed in GABAergic and glutamatergic neurons and there is significant correlation for its Gene Significance with intramodular connectivity and in module-trait relationship for this module and its associated cell types. Our analysis further showed distinct differences in function of different cell types between mice of opposing levels of stress-vulnerability. The PFC of mice which were more likely vulnerable to stress, as reflected by their high *EF*1 score, were enriched in processes such as cellular respiration, ribosomal function, mitochondria activity, and microglia activation. In contrast, mice which demonstrate less stress vulnerability brain activity (i.e. lower *EF*1 scores) have cell transcriptional states involved in synaptic processes for neuronal activity across all cell types.

While the number of studies that have broached studying this latent vulnerability state is small, there are several studies of note in the context of which we interpret our findings. These studies include identification of a stress vulnerability state via behavioral, neural circuit, and peripheral identifiers ^45–48^. Encouragingly, our findings are largely consistent with the findings in these studies. For example, a previous study showed that IL-6 levels from peripheral blood or by ex vivo stimulation of leukocytes could predict susceptibility or resilience to a subsequent stressor. By replacing the peripheral immune system in stress-naïve mice with those from stressed animals, this could increase their susceptibility to repeated social defeat stress ^45^. While this study addresses induction of vulnerability peripherally and our study measures gene expression in the PFC, our data are consistent with this picture of increased immune response in the latent stress vulnerable state. We observed gene expression consistent with increased activity of microglia, the cells of the brain resident immune system. We observed gene expression altered toward proinflammation and immune activation in mice with high *EF*1 activity. Many studies have shown that negative stressors can induce microglia activation (PMID: 31847911) which can rewire and impair neurocircuits ^49,50^ and is linked with various psychiatric disorders ^51,52^. To our knowledge, this is the first study to report elevation in microglia related function in the brain (and specifically PFC) of stress-naive vulnerable mice. However, regulation of microglia in PFC following chronic social defeat stress has been linked to stress susceptibility measured post-stress ^53^. Also, many studies of prenatal stress or early life adversity, which are often precursors to a vulnerable state, are consistent with our findings suggesting an important role for activated microglia in conferring vulnerability to stress ^54–57^.

In addition to microglia activation, our data also highlighted altered cellular respiration and mitochondria activity across all cell types in mice with high *EF*1 activity. While we had regressed out the influence of mitochondria percentage in our dataset from healthy cells and also subsequently removed all mitochondrial derived genes from our analysis, there are genes involve in mitochondria function that are derived from the nucleus ^58^. We not only observed enrichment for mitochondria related processes from genes more highly expressed in *EF*1 but also observed differential connectivity of genes involved in mitochondria related pathways in the pink module, which has the highest trait and DEG correlation across cell types. Furthermore, the top gene for intramodular connectivity and high Gene Significance in the pink module is involved in mitochondria related pathway which further suggests the importance of mitochondria function for the *EF*1 network.

Mitochondria provide the energy that fuels enzymatic reactions for neuronal processes, which can affect stress response and behavior ^59^. For example, a study where C57BL/6J mice were substituted with mitochondria containing a mutation that leads to the increased production of reactive oxygen species resulted in altered stress response, anxiety-like behavior and altered corticosterone regulation ^60^. Other studies have also linked mitochondria dysfunction to depression and other mood and psychiatric disorders where stress frequently plays an important role ^61–64^. Gene expression studies of major depressive disorder have link mitochondrial gene expression in the PFC with depression ^65^. In fact, a mitochondrial metabolite has been identified that significantly alters neuroplasticity in the PFC and hippocampus, with promising data suggesting its as a therapeutic target for depressive disorders ^66–68^. Follow up studies have found that factoring in mitochondrial and metabolic measures alongside early life stress experience can enhance prediction of antidepressant response ^69^.

Importantly, recent studies have linked mitochondria function to stress susceptibility ^70–72^. For example, mice exposed to chronic mild stress showed that susceptible and resilient mice had different expression of proteins involved in the mitochondria respiratory chain and mitophagy markers in the ventral hippocampus ^72^. Similarly, a genome-wide study of gene expression in the PFC and NAc revealed that mice exposed to multimodal chronic stress had altered expression of mitochondria genes in the PFC of susceptible mice and the mitochondria gene expression profile correlated with behavior in these mice ^73^. Given that these studies highlight the role of mitochondria in regulating behavior, our finding that respiratory and mitochondria processes are being upregulated in mice with high *EF*1 activity suggests that altered mitochondria function could contribute to vulnerability to stress prior to stress exposure.

Given that our differential gene expression profile is connected to stress vulnerability, we wanted to evaluate how this would correspond with gene expression profiles of mice that were susceptible to stress after stress exposure. Although we saw a few overlapping genes between our DEG data set and those of Bagot et al ^9^, one possible explanation for this could be due to comparison our single-cell resolution findings to data collected from bulk RNAseq dataset. In line with this, we saw overlaps in DEGs from the two cell types (glutamatergic neurons and astrocytes) with the most numbers of cells. Interestingly, we saw many more overlapping genes (hundreds vs tens) when we compared our DEGs against the broader networks of susceptibility and resilience in Bagot, R. et al and of the hub genes identified as key drivers of susceptibility, we find overlapping genes across all cell types. This provides a narrow list of key genes to test with regard to regulation of stress vulnerability in specific cell types moving forward. It is possible that the differences and similarities from the two gene sets are biologically significant, comparing the latent vulnerability state in animals that have not experienced chronic stress to those who have. No doubt there are many compensatory changes that happen in response to stress that manifest as a depressive state. It is possible that the overlapping genes between our two datasets point to genes that are pre-existing causal genes in the development of a depressive phenotype (Table S7, Summary_Overlap_with_Bagot, column C-D, cell-type specific DEG, column J). Interestingly, we observed little to no overlap when comparing with mice that had a recent stress exposure. However, we did observe more overlapping genes at a time further from the stress exposure (28 days post chronic social defeat). From these overlapping genes, we observed enrichment for ribosomal processes, which is consistent with a previous study that observed dysfunction in ribosomal gene expression in mice exposed to chronic social defeat stress in other brain regions ^74^. Overall, these finding largely suggest that genes involved in stress vulnerability could be distinct from those altered by stress in susceptible animals.

In summary, this is the first study that has attempted to identify cells, transcriptional architecture, and biological processes involved in latent stress vulnerability in the PFC, an important regulatory hub in the brain involved in the development of stress-susceptible phenotypes. To do this, we used a novel approach of leveraging electrical brain-wide network activity (i.e. elevated *EF*1 activity) from multi-site in vivo recordings combined with single cell RNA-Seq to start to unravel the complex interplay of different cell types in mediating this brain state. While our study suggests that all major cell types are involved in the process, it highlights the importance of GABAergic neurons in this process and also sheds light on how microglia and mitochondrial activity could be involved in the development and maintenance of this brain state.

## Acknowledgements

This work was supported by a NARSAD Young Investigator Grant from the Brain & Behavior Research Foundation (Rainbo Hultman). Additionally, this project received funding by *1DP2MH126377-01 (Rainbo Hultman),* the Roy J. Carver Charitable Trust (Rainbo Hultman), and NINDS T32NS007124 (Yassine Filali and Micah Johnson). We would like to thank Jens Hjerling-Leffler, Hayley French, Pat Sullivan, and Shuyang Yao for shared technical expertise. We thank Julia Miller for early technical support, and Neil Gallagher, Ted Abel, and Thomas Nickl-Jockschat for helpful conversations. We thank the Iowa Institute of Human Genetics Genomics Division for training and use of equipment.

## Author Contributions

Conceptualization, RH and BH, Methodology RH, YF, WW, ME, BH, SBM, Validation BH, ME and SBM, Formal Analysis, BH, IH, SS, MM, Investigation BH, SBM, ME, Writing Original Draft RH, BH, ME, SBM, IH Writing Reviewing & Editing RH, BH, SBM, ME, YF, MM, IH, SS, Visualization MM, BH, SBM, IH, Supervision, RH, Project Administration, RH, BH, SBM, Funding acquisition, RH.

## STAR Methods

### Lead contact

Further information and requests for resources and reagents should be directed to and will be fulfilled by the lead contact, Rainbo Hultman

### Materials availability

This study did not generate new unique reagents.

### Data and code availability

Single-cell RNA-seq data will be deposited at Gene Expression Omnibus (https://www.ncbi.nlm.nih.gov/geo/) and be made publicly available as of the date of publication. Accession numbers are listed in the key resources table. Neurophysiological data reported in this paper will be shared by the lead contact upon request.

Adaptation and parameters of the standard available R packages are available in the STAR Methods section. Statistical analysis of cell-cell interaction analyses has been adapted from [] as stated in the STAR Methods section.

Codebase for GAM analysis and producing the *Electome Factor* scores can be found at: [link for review process (will be made publicly available upon publication) https://gitlab.com/rainbo-hultman/electome-factor-1.git?private_token=glpat-sWnRGsUtVh-vfvSo16Nt]

Any additional information required to reanalyze the data reported in this paper is available from the lead contact upon request.

### EXPERIMENTAL MODEL AND SUBJECT DETAILS

#### Animals and Experimental Design

All animals were housed in accordance with the University of Iowa Institutional Animal Care and Use Committee (IACUC) standards. All mice were housed in bedded cages and exposed to a 12-hour light/dark cycle with ad libitum access to food and water. This study utilized C57BL/6J mice delivered from the Jackson Laboratory in Bar Harbor, Maine at 5-7 weeks of age. Prior to surgery, experimental mice were group-housed by sex. Following surgical implantation of recording electrodes, experimental mice were singly-housed and placed on corncob bedding. Each implanted animal was exposed to a forced interaction test (FIT), a test designed to elicit a negative affect response, one time only: two days prior to sample collection for single-cell RNA sequencing (scRNA-seq) (Fig 1). Forty-eight hours after recording each FIT, the prefrontal cortex (PFC) was micro-dissected and processed for scRNA-seq.

#### Surgical Implantation of Electrodes

For surgical implantation of recording electrodes, mice were anesthetized with 1-5% isoflurane, and their skulls were secured in a stereotaxic device. Metal ground screws were placed above the anterior cranium and the cerebellum before implanting target bundles. Electrode bundles designed to target the following regions: nucleus accumbens (NAc), amygdala (AMY, basolateral, BLA and central, CeA), ventral hippocampus (VHipp), and ventral tegmental area (VTA), were positioned according to stereotaxic coordinates measured from Bregma as previously described ^75,76^. These coordinates were chosen to maintain consistency with prior molecular and neurophysiology studies of vulnerability following chronic stress ^9,21^. The PFC (including prelimbic, Prl and infralimbic, IL components) was intentionally left out of the electrode design so as not to disturb the tissue that would later be collected for scRNA-sequencing. This was possible because *EF1* scores can still be calculated with minimal regions missing from the recording data ^21^. Experimental animals were implanted between 6-10 weeks of age and were allowed to recover for a minimum period of 2-weeks prior to testing (Figure S1). Accuracy of electrode placement during implantation was evaluated at the conclusion of the experiments through individual subject histological assessment. This was done to confirm the exact recording sites used to capture neurophysiological data.

#### Acquisition of Neurophysiological data

Neurophysiological data was captured as previously described ^21^ for each subject during the completion of a forced interaction test (detailed below). The CerePlex Direct acquisition system was used to capture neuronal activity data sampled at 30kHz (Blackrock Microsystems, UT, USA). Acquired local field potentials (LFPs) were stored at 1000 Hz following a bandpass filter set at 0.5-250Hz. A ground wire connecting both ground screws served as an overall reference for neurophysiological recordings within each experimental animal.

#### Forced Interaction Test

The forced interaction test (FIT) was performed as previously described ^77^ with recordings obtained from 5 brain regions over the duration of the FIT in addition to the capture of corresponding video for animal behavior (Figure 1A) ^21^. Mice were first connected to a mu headstage akevlarlar-coated cable (Blackrock Microsystems, UT, USA) before being placed in a fresh homecage with corncob bedding. Following a 30- minute habituation period, a 10-minute baseline homecage recording portion of the FIT was performed. An 18” high, clear, acrylic cage extender was placed just inside the walls of the home cage to prevent the implanted animal from climbing out. Succeeding the homecage recording, the experimental animals were placed in a 3.25” diameter x 7” high covered Plexiglass carousel (Noldus) while neurophysiological data was collected for a period of 5 minutes. A CD1 mouse previously screened for aggressive tendencies was then placed inside the cage extender, but outside the carousel within the homecage, and the recording continued for an additional 5 minutes. The carousel was designed to protect experimental animals from outside harm, while maintaining sensory interactions with the outside environment through the presence of open-air slots. Experimental mice were only subjected to the FIT once, a minimum of 2 weeks following surgical implantation of electrodes to allow for sufficient recovery, and always 2 days prior to tissue collection for scRNA-seq sample prep. FIT recordings and subsequent sample collections were performed when animals were between 9 and 14 weeks of age (Figure S1). The local field potentials (LFPs) obtained from the FIT recordings were evaluated for each mouse to determine their *Electome Factor 1* (*EF1*) scores, a previously identified marker of the stress vulnerable state (see below).

#### Buffer Preparation, Tissue Collection, and Generation of Single-Cell Suspensions

Artificial cerebrospinal fluid (NMDG-aCSF) buffer was freshly prepared on the day of harvesting cells and contained the following powdered reagents, in order: 96 mM NMDG (Sigma-Aldrich; St. Louis, MO; CAS No. 6284-40-8,), 2.5 mM KCl (Research Products International; Mt. Prospect, IL; CAS No. 7447-40-7), 1.35 mM NaH_2_PO_4_ (Sigma-Aldrich; St. Louis, MO; CAS No. 7558-80-7), 30 mM NaHCO_3_ (Research Products International; Mt. Prospect, IL; CAS No. 144-55-8), 20 mM HEPES (Research Products International; Mt. Prospect, IL; CAS No. 7365-45-9), 25 mM glucose (Research Products International; Mt. Prospect, IL; CAS No. 50-99-7), 5 mM Na+ascorbate (Sigma-Aldrich; St. Louis, MO; CAS No. 134-03- 2), 2mM thiourea (Sigma-Aldrich; St. Louis, MO; CAS No. 62-56-6), 3 mM Na+pyruvate (Sigma-Aldrich; St. Louis, MO; CAS No. 113-24-6), 2 mM N-acetyl-L-cysteine (Sigma-Aldrich; St. Louis, MO; CAS No. 616-91- 1). Powdered reagents were measured using weigh boats and poured into a 500mL graduated cylinder one at a time; excess material was washed into the graduated cylinder using ddH20. Following the addition of all powdered reagents, the solution was then adjusted to 80% final volume with ddH20. Under the fume hood, the liquid reagents were then added directly to the solution using a P1000 pipette, in the following order: 37% HCl (Sigma-Aldrich; St. Louis, MO; CAS No. 7647-01-0), 0.5 mM CaCl2 (Alfa Aesar; Ward Hill, MA; CAS No. 10043-52-4), 10 mM MgSO4 (GBiosciences; St. Louis, MO; Cat. # 786-530). The final volume of the solution was then adjusted to 500mL using the ddH20 spray bottle. After adding a stir bar into the solution and covering it with parafilm, the graduated cylinder was placed atop a stir plate, and all reagents were allowed to dissolve into the solution at 300-350 rpm for 10 minutes. The incorporated NMDG-aCSF solution was then filtered through a 100µm Nylon cell strainer (Falcon; Corning, NY; Ref: 352360) into an Erlenmeyer flask, covered with parafilm, labeled, and either stored at 4°C overnight or prepared for use as follows.

NMDG-aCSF, prepared within 24-hours of planned collections, was bubbled in carbogen (95% O2/5% CO2) for at least 75 minutes on ice prior to contacting any tissue samples. During this time, Earle’s Balanced Salt Solution (EBSS) solution and albumin inhibitor solution were also bubbled on ice, which was maintained throughout the duration of the experiment, taking care to prevent the formation of bubbles in the albumin inhibitor. After the NMDG-aCSF had been bubbling for 75-100 minutes, the pH was measured to be 7.35-7.45 and the osmolarity was measured to be 300-310 mOsm. If pH or osmolarity measurements fell outside of their respective range, then the buffer was not used for sample processing.

Animals were anesthetized using Euthasol (Virbac AH, Inc.; Fort Worth, TX) and perfused with ice cold NMDG-aCSF. Their brains were rapidly removed, adhered to a rotating specimen disk using Loctite® 404™ Quick Set™ Instant Adhesive (Henkel; Rocky Hill, CT; P/N 135465), placed in a cold bath of bubbling NMDG-aCSF buffer on a Leica VT1200 vibratome (Leica Biosystems; Ref: 14048142065) and sliced coronally into 200µm slices. Exactly six slices were used for each brain’s PFC dissection (Figure S3). Ice was used to keep the samples cold whenever possible in all proceeding steps.

The slices were transferred to recovery holding strainers secured in 6-well plates and bubbled in NMDG-aCSF. Slices were allowed to recover for 10-15 minutes. Papain solution was prepared by adding oxygenated cutting solution to the papain and incubating this without agitation at 37°C for 10 minutes. Five minutes into the papain incubation, oxygenated EBSS was added to the DNASE, mixed by inversion, and placed on ice. After 10 min of papain incubation, the papain was combined with the prepared DNASE, mixed by inversion, and left at room temperature. While the papain was being prepared, the slices were micro-dissected (Figure S3) using a scalpel and dissection scope (Jewelmont) with a halogen cool light source (AmScope; SKU: HL250-AY; Irvine, CA) and transferred to a microcentrifuge tube. The NMDG-aCSF was then carefully removed from the tube containing the micro-dissected tissue, being replaced by the addition of the papain solution. The tissue was incubated in the papain solution for 30- 40 minutes at 37°C at 60 RPM using an Innova 4000 Incubator Shaker (New Brunswick Scientific; Enfield, CT; P/N: NB-4000). The papain solution was removed from the tissue and diluted albumin inhibitor (stop solution) was added. Triturations were carried out using two sequential bore diameters (∼300µm and ∼150µm) of fire-polished glass capillaries (World Precision Instruments; Sarasota, FL, Item No. TW100-3). A total of six triturations were used for female mice and twelve triturations for male mice, with the first half completed using the 300µm bore capillary and the latter half with the 150µm bore capillary. After the mechanical dissociation was complete, the resulting suspension was cloudy and homogenous. The suspension was filtered through a 40um mesh filter (Falcon, Corning NY; Ref: 352340), and then transferred to a 15mL conical tube and centrifuged at room temp., 0.1 RCF for six minutes. Following centrifugation, the supernatant was quickly but carefully removed. Using a P1000 wide-bore tip, the pellet was resuspended in 1mL of stop solution (aspirated and suspended about five times) and then filtered through another 40µm mesh Falcon cell strainer. In another conical tube, 5mL of stop solution was added, and the cell suspension was carefully layered on top using a P1000 wide-bore tip. This suspension was again centrifuged at room temp, 0.1 RCF for six minutes. Following the second centrifugation, a P1000 wide-bore tip was used to remove the supernatant until about 50µL of the solution was left. Then, using a P200 tip, 20-30µL of the solution was removed until about 20µL of cell suspension remained. The pellet was then resuspended in 30µL HBSS to generate the final single-cell suspension.

#### Cell Counting

Equal volumes of single-cell suspension and 0.4% trypan blue stain (Life Technologies corporation; Grand Island, NY; Ref: 15250-061) were mixed and 10µm of the resulting solution was loaded onto a hemacytometer (Hausser Scientific; Horsham, PA; Cat No. 02-671-10). A Leica DMi1 inverted microscope (Leica Biosystems; Ref: 11526227) was used to view the counting chamber using an S40-slider (Leica Biosystems; Ref: 1152613). Both live and dead cell counts were obtained from each of the four corners of the chamber’s etched grid to determine total cell density as well as the percentage and total of live cells present in the single-cell suspension.

#### Projecting Neurophysiological Data into ‘Electome Factor Space’

LFP data acquired from the FIT recordings were used in conjunction with the learned feature representation from a previously developed machine learning model using “discriminative cross-spectral factor analysis” (dCSFA) to calculate vulnerability scores (*Electome Factor* 1) ^21^. The LFP oscillations obtained from our experimental mice were projected into the space of the *Electome Factors* that were previously learned as previously described ^21^.

#### Single cell library preparation and sequencing

Single cell libraries were generated using the Chromium Next GEM Single Cell 3’ Kit v3.1 as per manufacturer’s instructions and sequenced using the Illumina NovaSeq6000 using 100 bp paired-end reads at the University of Iowa Institute of Human Genetics Genomics Division. Raw base call files were demultiplexed and FASTQ files were generated using the Cell Ranger program (10X Genomics).

#### Quality control, clustering and celltype identification

FASTQ files (N=168) were evaluated using FastQC (v0.11.7) and MultiQC (v1.5) to determine excellent sequencing quality (Phred score > Q30). The cellranger count function from Cellranger-6.0.0 was used to align the FASTQ files to the mouse reference genome (mm10; pre-built reference mm10-2020-A obtained from the 10X genomics website) and for feature quantification. The average number of reads per cell was ∼ 97,000 reads (Table S2). Cellranger aggr was performed with the argument ‘--normalize mapped ’ to aggregate the data. Data was subsequently loaded into RStudio (4.1.1) and processed using Seurat (4.1.1) for normalization, data reduction, clustering, and visualization. Briefly, high quality cells defined by percentage mitochondria < 10% ^78^ and total feature counts > 1000 were retained for downstream analysis. The SCTransform function was used to normalize the data and control for confounding sources of variation such as read depth and mitochondria percentage. Based on knee plots, 15 principal components were used to create a shared nearest neighbor graph. Cells were clustered into groups by optimizing a modularity function (Louvain algorithm, resolution 0.8, 10 random starts and 10 iterations). Data was visualized using t-distributed stochastic neighbor embedding (tSNE) plots. Doublets were identified using scDblFinder (1.8.0) ^79^ and removed from the analysis. The number of doublets was ∼ 2% which is consistent with the expected number of doublets for 10X single cell RNA-Seq as per manufacturer’s guidelines. Cluster identity was determined by cell type specific marker gene expression including glutamatergic neurons (*Slc17a7* and *Slc17a6*), GABAergic neurons (*Gad1* and *Gad2)*, astrocytes (*Aldoc* and *Apq4)*, mature oligodendrocytes (*Mbp* and *Cldn11*), oligodendrocyte precursors (*Pdgfra),* endothelia (*Pecam1* and *Slc2a1)*, microglia (*Tmem119)*, pericytes (*Vtn)*, endothelia (*Pecam1* and *Slc2a1*) and vascular smooth muscle (*Tagln)*. To further validate the identity of the cells in each cluster, the cell2cluster function from the scmap package (version 1.16.0) was used together with a prefrontal cortex (PL-ILA-ORB) reference dataset extracted from the 10X single cell RNAseq dataset from the Allen Brain Atlas ^80^. Cells that were not consistent with the identity of their cluster were removed from the analysis. Similarly, a cluster that ambiguously co-expressed markers for endothelial and pericytes was also removed from the analysis.

#### Hierarchical clustering

*Electome Factor* 1 (*EF*1) score was used to calculate the Euclidean distance followed by hierarchical clustering using the complete-linkage method.

#### Differential gene expression

The differential gene expression analysis of the scRNA-seq results was limited to those animals having either high or low EF1 scores, excluding those with intermediate scores (Figure 1D). Only cell types with at least 10 cells per individual mouse were used for differential gene expression (DEG) analysis using corrected counts obtained from SCTransform as a recent study showed that this reduces false positive rates ^81^. All mitochondria-transcribed genes were removed prior to DEG analysis. Analysis was performed using ZINB-WaVE-DEseq2 workflow ^82^. Size factors were computed using computeSumFactors ^83^. Significance testing was performed using the likelihood ratio test to evaluate the effects of EF1 while controlling for sex as a covariate in the full model (∼ EF1_level + Sex) and compared against the reduced model (∼ Sex). Additional arguments included in the Deseq function were useT=TRUE, minmu=1e-6, minRep=Inf and sfType=“poscounts” as per recommendation in the DESeq2 and ZINB-WaVE vignettes and other studies ^24,82,84,85^. In order to assess the validity of the SINB-WaVE differential gene expression, the package DESingle ^27^ was also used for DEG analysis as it is comparatively as good or better in performance compared to other algorithms ^86^ and we have successfully used this method for DEG analysis in a previous study ^87^. Genes with FDR < 0.05 were differentially expressed.

#### Gene Ontology analysis

Gene ontology analysis was performed using clusterProfiler (version 4.2.2) ^88^ or EnrichR (version 3.2) ^89^ to identify biological processes that were over-represented for genes that were differentially expressed. We also used SynGO ^28^ to identify over-representation of genes for biological processes and cellular components related to the synapse.

#### Gene Set Enrichment Analysis

Normalized gene expression counts from DESeq2 was used for gene set enrichment analysis (GSEA). Analysis was performed using GSEA (version 4.2.1) with the gene ontology biological processes dataset (C5) containing 7751 gene sets from the Molecular Signatures Database or using mitochondria reference dataset from the MitoCarta 3.0 ^26^. Mouse gene symbols were remapped to their human orthologs using GSEA program and we also used the following settings: phenotype for permutation type, enrichment statistic was classic and metric for ranking genes was Signal2Noise.

#### Comparison of Differentially Expressed Genes with Genome-Wide Association Study (GWAS) datasets

Magma (v1.10) was used to compare differentially expressed genes from high and low EF1 groups with GWAS datasets from the Psychiatric Genomics Consortium. These datasets include Attention Deficit Hyperactivity Disorder^90^, Alzheimer’s Disease ^91^, Anxiety Disorder ^92^, Autism Spectrum Disorder ^93^, Bipolar Disorder ^94^, Anorexia Nervosa ^95^, Major Depressive Disorder ^96^, Obsessive Compulsive Disorder ^97^, Tourette Syndrome ^98^, Post Traumatic Stress Disorder ^99^, and Schizophrenia ^100^. Magma analysis was performed as previously described ^101^. P-values were corrected for multiple testing using false-discovery rate.

#### Upstream Regulator Assessment via Ingenuity Pathway Analysis (IPA)

We examined potential upstream regulators as predicted by our differential gene expression by cell type. Qiagen’s Ingenuity Pathway Analysis (IPA), a unified, web-based application that enables integrated analysis and elevated understanding of such complex data sets was used for this purpose. This software is supported by the Ingenuity Knowledge Base, which is composed of biological and chemical findings of extreme detail, allowing for targeted searches and improved understanding of a dataset’s significance within a broader biological context (Qiagen, 2023).

#### Network analysis using hdWGCNA

High Dimensional Weighed Gene Correlation Network Analysis (hdWGCNA) was performed using the R package as described in the vignette ^102^. Briefly, genes that are expressed in at least 5% of cells were retained for the analysis. To overcome data sparsity inherent to single cell data, a metacell gene expression matrix was created using the function “MetaCellByGroups” which aggregates SCTransform count data from cells of each cell type and from the same biological sample of origin. Only cell types that met the default setting of 100 cells for each sample were retained for the analysis (i.e. GABAergic neurons, glutamatergic neurons, astrocytes and microglia). Data was then log normalized using the “NormalizeMetacells” function and expression matrix of all cells types were specified for network analysis using the “SetDatExpr” function. Using the “TestSoftPowers“ function, a soft-power threshold was calculated according to the scale-free topology criterion ^30^ for a scale-free topology index (R^2^) of > 0.8. The “ConstructNetwork“ function was subsequently used to calculate the adjacency matrix, topological overlap matrix (TOM), and the Dynamic Tree Cut algorithm to identify gene modules. Data when then scaled and centered using “ScaleData” function followed by calculation of module eigengenes for single cells using the “ModuleEigengenes” function. Eigengene based connectivity was calculated using the “ModuleConnectivity” function. The relationship between modules and EF1 groups was assessed by Pearson correlation between module eigengenes for each cell type and EF1 groups using the “GetModuleTraitCorrelation” function. The “FindDMEs” function was used to evaluate how eigengenes expression varies between EF1 high and low groups for each cell type. To determine enrichment of differentially expressed genes in modules, the “OverlapModulesDEGs” function was used which uses the Fisher’s Exact Test to assess for over-representation of DEGs in modules. Gene Significance was calculated as previously described by taking the absolute value of Pearson correlation between metacell normalized gene expression and EF1 group ^30^.

#### Differential Network Analysis

Metacell normalized gene expression matrix for each cell type was used to evaluate differences in network structure within modules between high and low EF1 groups using the R package discoMod ^31^. This package uses p-norm difference (PND) test which improves sensitivity for detecting small proportion of module correlation changes between conditions. As PND to the exponent of 6 was previously shown to provide optimal performance ^103^, we therefore used this exponent for the analysis. To further evaluate how changes in network structure might affect gene regulatory pathways, the R package dnapath^32^ was used to determine pathways that have differences in network connectivity between EF1 high and low groups. The mouse pathways from the Reactome database and metacell normalized gene expression matrix for each cell type for each module ^104^ were used for this analysis.

