## Supplemental Materials for "Single Cell Transcriptome of Stress Vulnerability Network in mouse Prefrontal Cortex"

### Supplemental Experimental Methods

tSNE vs UMAP

To evaluate which clustering approach would be optimal for cell type identification, data was first clustered using tSNE and cluster identities were determined using cell type specific markers ( Figure S4). Data was then re-clustered using UMAP to determine if cluster identities would be maintained. Although cluster identities were broadly similar between tSNE and UMAP, cells that express *Slc17a7*, which is highly expressed in glutamatergic neurons, were clustered with microglia and GABAergic neurons in UMAP (Figure S4B, D). These cells were however clustered with glutamatergic neurons in tSNE (Figure S4A, C). Given that tSNE clustering grouped glutamatergic cells together as a cluster, we therefore used tSNE clustering for our analysis.

Evaluating cluster identity and cell type distribution

Cells were clustered by tSNE (Figure S5A) and cluster identities were determined using expression of cell type specific markers ( Figure S5B-O). As the Mouse V1 and Alm SMART-Seq single cell datasets from the Allen Brain Atlas had designated clusters expressing both endothelia and pericyte markers as low-quality doublets, we therefore remove that cluster from further analysis. The distribution of the remaining cell types was similar between males and females (Figure 2C). Interestingly, the percentage of GABAergic neurons was significantly elevated in the high EF1 group (FDR < 0.05) (Figure 2D).

Evaluating the effect of covariates on tSNE clustering

We evaluated how cell cluster identities (Figure S5A) might be affected by different covariates. Overlapping covariates such as sex, EF level and experiment date with tSNE clusters showed homogeneous distribution suggesting that these factors have little influence over cell clustering

(Figure S6B-D).

Categorization of GABAergic neurons

To determine subtypes of GABAergic neurons in our dataset, GABAergic neurons were sub-clustered and their cell identities assigned by the cell2cluster program [1] using a reference mouse prefrontal cortex 10X dataset [2] (Figure S7A). We identified cells that had low or no *Gad1* or *Gad2* expression (Figure S6B) and were not assigned any cell type by cell2cluster. As such, these cells were removed from further analysis. We also identified previously published subtypes of GABAergic neurons expressing *Sst*, *Pvalb*, *Chodl*, *Lamp5*, *Vip* and *Sncg* (Figure S7C-H).

Distribution of GABAergic neuron subtypes and differential gene expression

The distribution of the subtypes is similar to that of a previous study [2] but with ~10% more *Pvlab* and ~10% less Vip expressing cells (Figure S8A). Although there was a higher percentage of *Pvalb* expressing cells in *EF*1 high group compared to *EF*1 low group and a lower percentage of *Vip* expressing cells in *EF*1 high group compared to *EF*1 low group, these observations did not reach a significance threshold padj<.05 (Figure S8B). Using ZINB-WaVE + DESeq2 , we also performed differential gene expression for GABAergic neuronal subtypes that had at least 10 cells for all samples in our dataset. We observed significantly more DEGs for *Sst* expressing neurons compared to *Pvalb* expressing neurons (Table S11). For both subtypes, there were more genes that were more highly expressed in the high EF1 group (Table S11).

Differential genes expression using DESingle

To validate our differentially expressed genes from ZINB-WaVE + DESeq2 analysis, we performed another independent analysis using DESingle for all the cell types which has a minimum of 10 cells for all samples in our dataset (Figure S9). We further performed Gene Ontology analysis to determine biological processes that were over-represented with differentially expressed genes from each cell type (Figure S10). For the high *EF*1 group, top biological processes pertained to respiration and mitochondria processes for glutamatergic and GABAergic neurons and astrocytes (Figure S10A-C). Microglia and astrocytes were enriched for biological processes involved in synapse related processes (Figure S10D-E). In contrast, all cell types for the low *EF*1 group were enriched for synapse related processes (Figure S10F-J).

Analysis of differentially expressed genes by SynGO

Since our previous analysis showed that synaptic related processes were enriched in the low *EF*1 group compared to the high *EF*1 group, we validated this finding by performing an over-representation analysis using SynGO. Consistent with our previous analysis, genes which were more highly expressed in the high *EF*1 group had little to no over-representation in synapse related processes (Figure S11A-E). By comparison, the low *EF*1 group had consistently more over-representation in synapse related processes for the respective cell types (Figure S11F-J).

Electome Factor 1 Score Distributions

In order to determine the nature of our experimental mouse *EF*1 scores within the range of all possible *EF*1 scores, we plotted the distribution of *EF*1 scores from 85 mice, including data from our previous paper [3] (Figure S12). Given that the data are skewed to the right, we then applied three different transforms of the data (Figure S13). All of these distributions are bimodal, but the fourth-root transformation appears to have a mixture distribution with nearly a point mass close to zero and a normal distribution with a mean close to 0.5. Using the mixtools R package, we estimated the fourth-root transformed *EF*1 scores to have a mixture normal distribution with means: $\mu_{1}=$ 0.0820851, $\mu_{2}=$ 0.4062096 and standard deviations: $\sigma_{1}=$ 2*.*5604307 × 10^−7^, $\sigma_{2}=$ 0.1647875 [5]. The square-root and log transformed densities appear to be bimodal with a skewed second component density; therefore, we used the fourth-root transformation for further examination of the relationship between cell types and *EF*1 scores.

Generalized Additive Model

GAMs were used to capture additive non-linear relationships between the response and predictors with regard to principal components for each cell type and the fourth-root transformed EF1 scores. Given that the number of samples (mice) is small compared to the number of gene expression values for each cell type, we separately modeled the effects of principal components for a single cell type on the fourth-root transformed EF1 scores. Specifically, if $y_{i}$ and $x_{i}$ are the fourth root of *EF*1 score and first or second PC score for mouse $i$, respectively, then GAM posits that

$$y_{i}=f\left( x_{i} \right)+\epsilon_{i}, i=1, \ldots, 11, E\left( \epsilon_{i} \right)=0, \text{var}(\epsilon_{i})<\infty,$$

where $f$is the unknown regression function and is twice continuously differentiable. We fit these models using the mgcv R package. The tuning parameter for choosing the function smoothness in the mgcv package is selected using generalized cross-validation [4].

### Supplemental Results

Determining Relationships of Cell type with Electome Factor 1 Score

We used single cell count data and *EF*1 scores for all twelve mice to determine cell-type/*EF*1 relationships. We applied fourth-root transformations to the *EF*1 scores based on the empirical distribution of previously computed *EF*1 scores and used those as the responses in our models. Figure S14 shows their mean *EF*1 scores, fourth-root transformed *EF*1 scores, and corresponding p-values under the individual normal distributions obtained from the mixture normal discussed above in the section “*Electome Factor* 1 Score Distributions”. Based on the p-values in Figure S14, the ninth mouse’s *EF*1 score could be from either part of the mixture components, so we chose to leave the ninth mouse out of our analysis and only use the other eleven mice whose *EF*1 scores almost certainly come from the upper part of the mixture normal distribution. For each of the remaining eleven mice in our analysis, we had single cell count data consisting of varying numbers of observations from five different cell types, each of which is a sparse vector providing the normalized cell counts associated with 17,549 different genes. An example of the normalized cell count data for a single mouse can be seen in Figure S15. For each mouse, we then averaged the normalized cell count vectors across each cell type. This resulted in a 5 × 17, 549 matrix for each mouse, an example of which can be seen in Figure S16. For each mouse’s 5 × 17, 549 mean cell count matrix, we performed principal component analysis (PCA) to reduce the dimensions of this matrix to 5 × 2. An example cell type by PC matrix can be seen in Figure S17. These combinations of cell type with principal component formed the covariates of our models.

We used generalized additive models (GAM) to estimate the effects of different cell types on the fourth-root transformed *EF*1 scores (see supplemental methods). GAMs capture additive non-linear relationships between the response and predictors. In our case, the number of samples (mice) is small compared to the number of gene expression values specific to the cell types; therefore, we separately modeled the effects of principal components for a single cell type on the fourth-root transformed *EF*1 scores. $y_{i}=f\left( x_{i} \right)+\epsilon_{i}, i=1, \ldots, 11, E\left( \epsilon_{i} \right)=0, \text{var}\left( \epsilon_{i} \right)<\infty,$

he plots in Figure S18 show the smoothing functions estimated for 10 GAM models, each fit to a single cell type with principal component combination, along with a two-standard error uncertainty region and the observed data used to fit the model. Table 2 shows for each cell type with principal component combination the adjusted R-squared and an approximate p-value for the null hypothesis that the effect of the smooth function of that cell type variable is zero. These results suggest that GABAergic cell types may produce a statistically significant, non-linear effect on the fourth-root transformed *EF*1 scores.

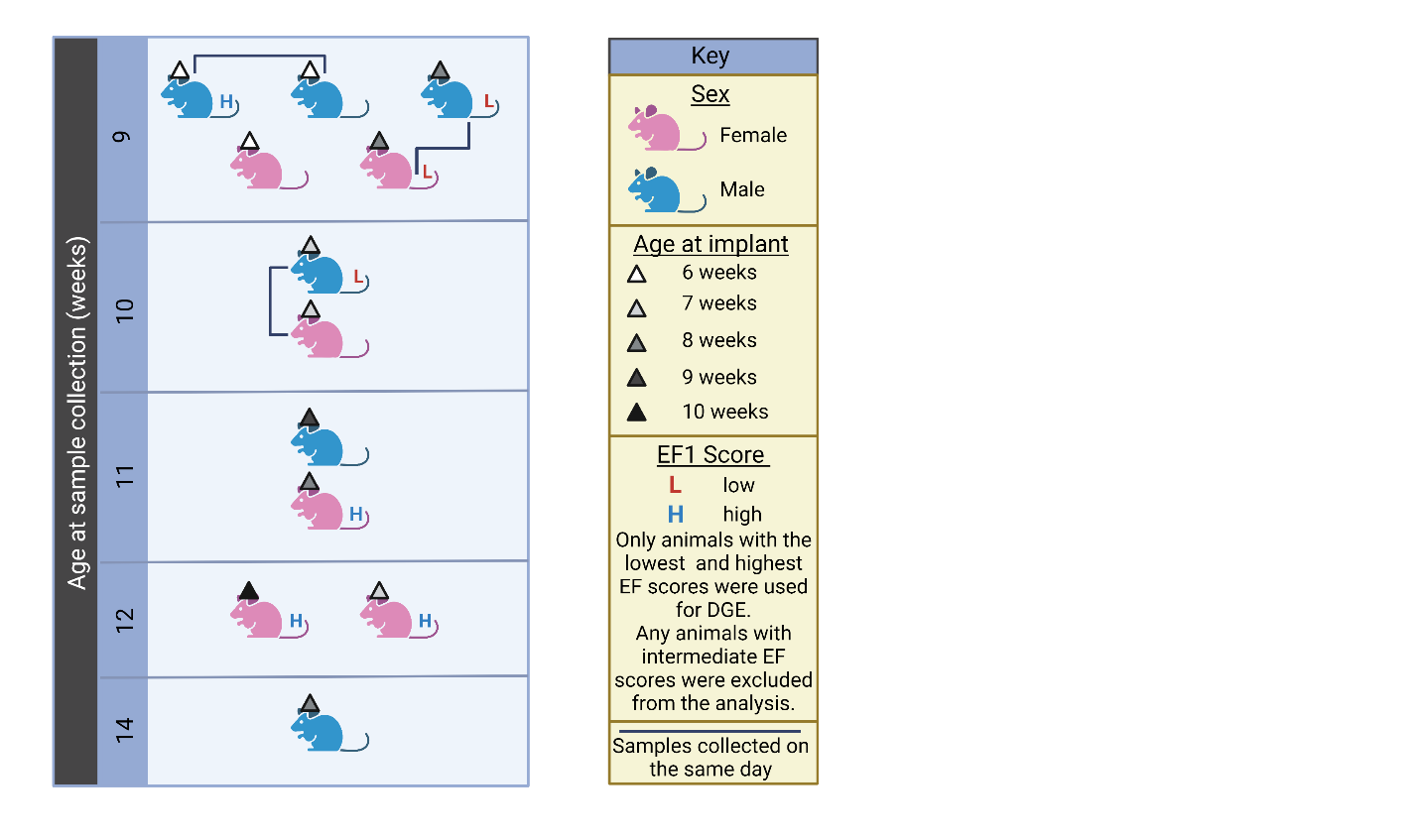

### Figure S1: Experimental sample collection timeline

Schematic diagram illustrating the age at which individual experimental animals were implanted and used for data and sample collections. Reference the key for appropriate age and sex color-coding information. Animals connected by a dark blue line had PFC samples collected and processed into single-cell suspensions on the same day. Only experimental animals with high or low *Electome Factor* 1 (*EF*1) scores were used in differential gene expression analyses.

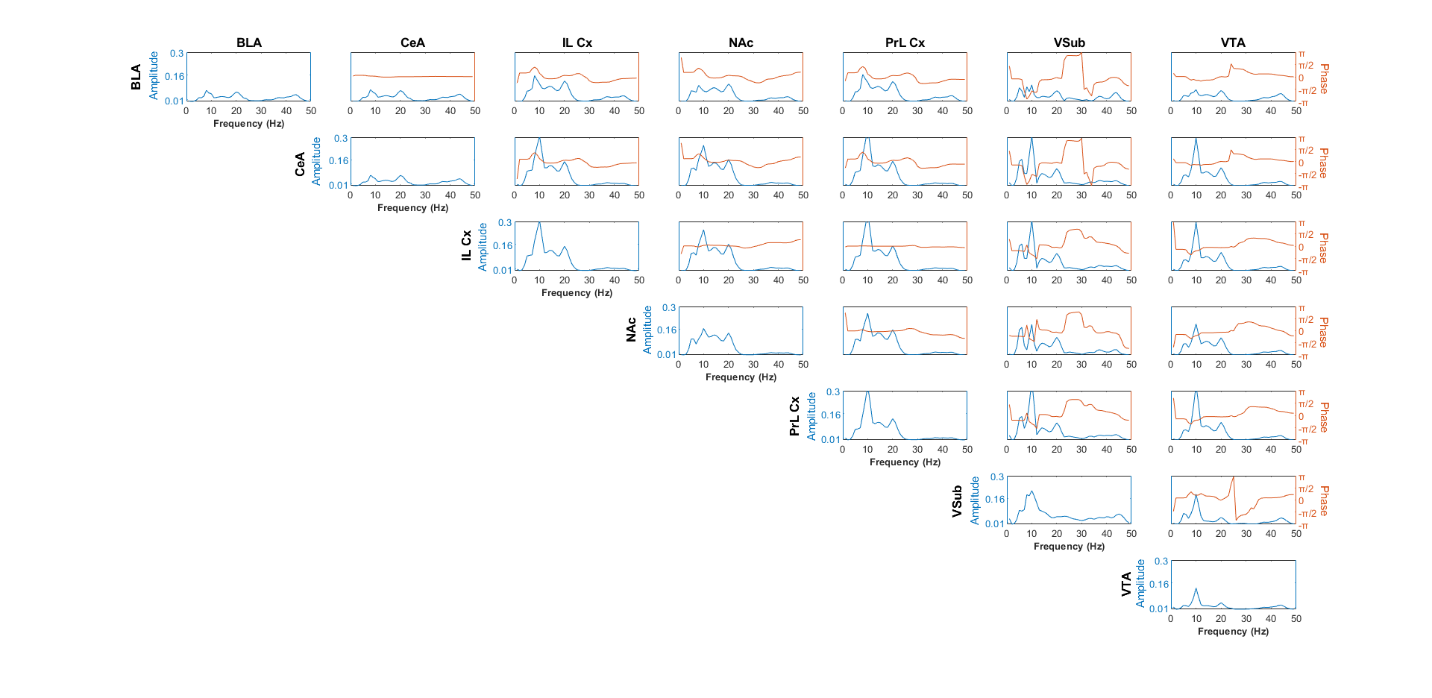

### Figure S2: *Electome Factor* 1 feature autocorrelation plots

Amplitude of power and coherence contributions to *EF*1 (left axes) are shown in blue. Phase/directionality contributions (right axes) are shown in orange. The x-axis for each plot represents frequency in Hertz (Hz).

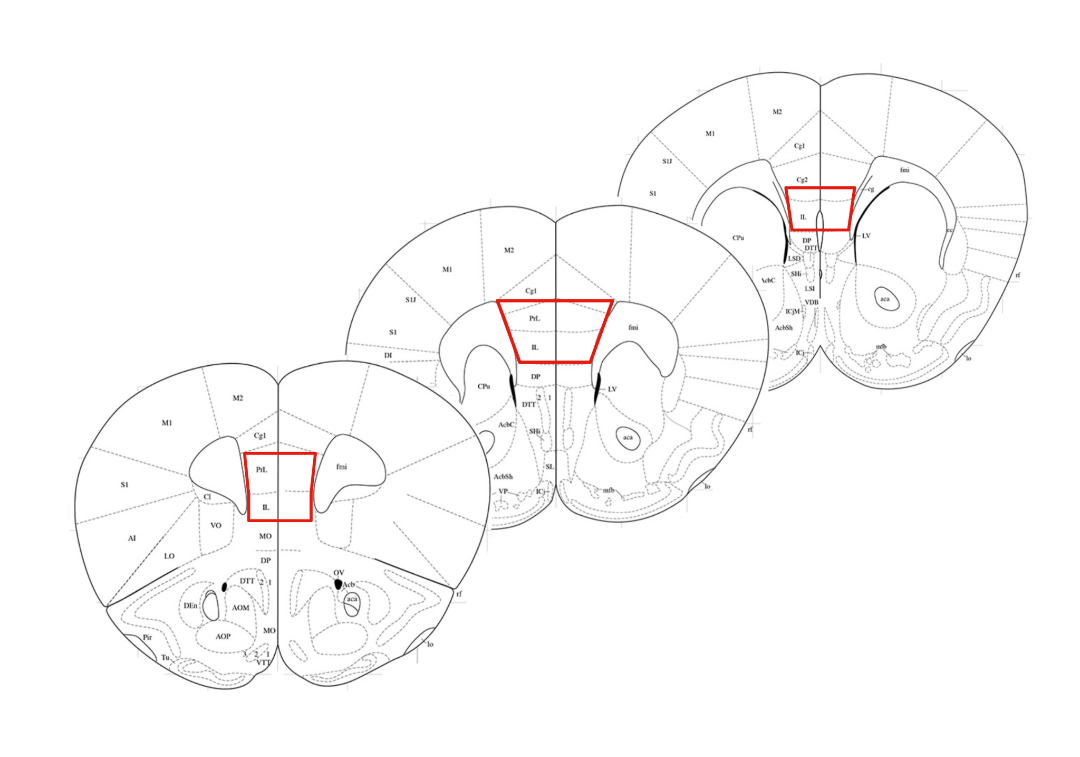

### Figure S3: Dissection Targets

Outlined in red lines are dissection cuts for prefrontal dissections. Each slice was 200 microns, and each dissection included exactly six slices.

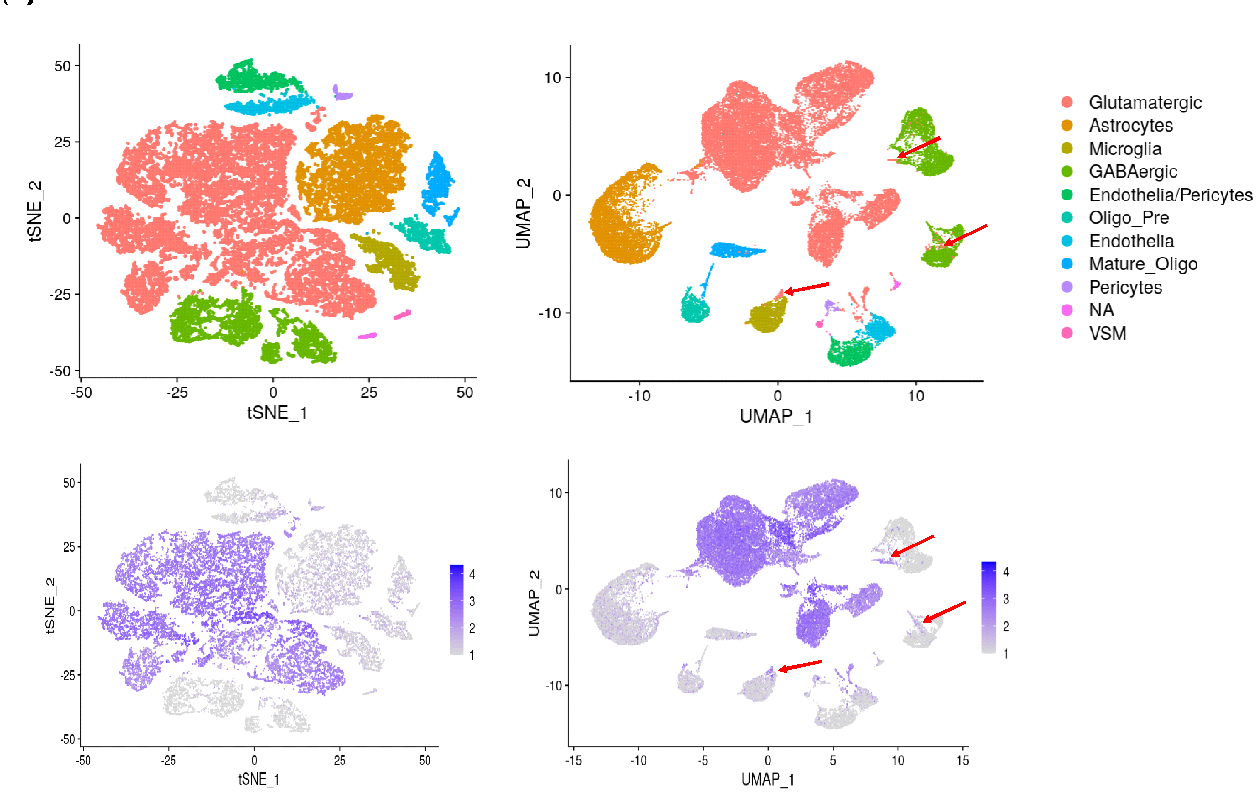

### Figure S4: Comparison of cell cluster identities by tSNE and UMAP

Data normalized by read depth from cellranger aggr was filtered to remove low quality cells. (left) Data was then clustered using tSNE and cell identities were determined using cell specific marker gene expression such as *Slc17a7* (bottom), a marker for glutamatergic neurons. (right) Data was then re-clustered by UMAP to determine if cells would be grouped similarly.

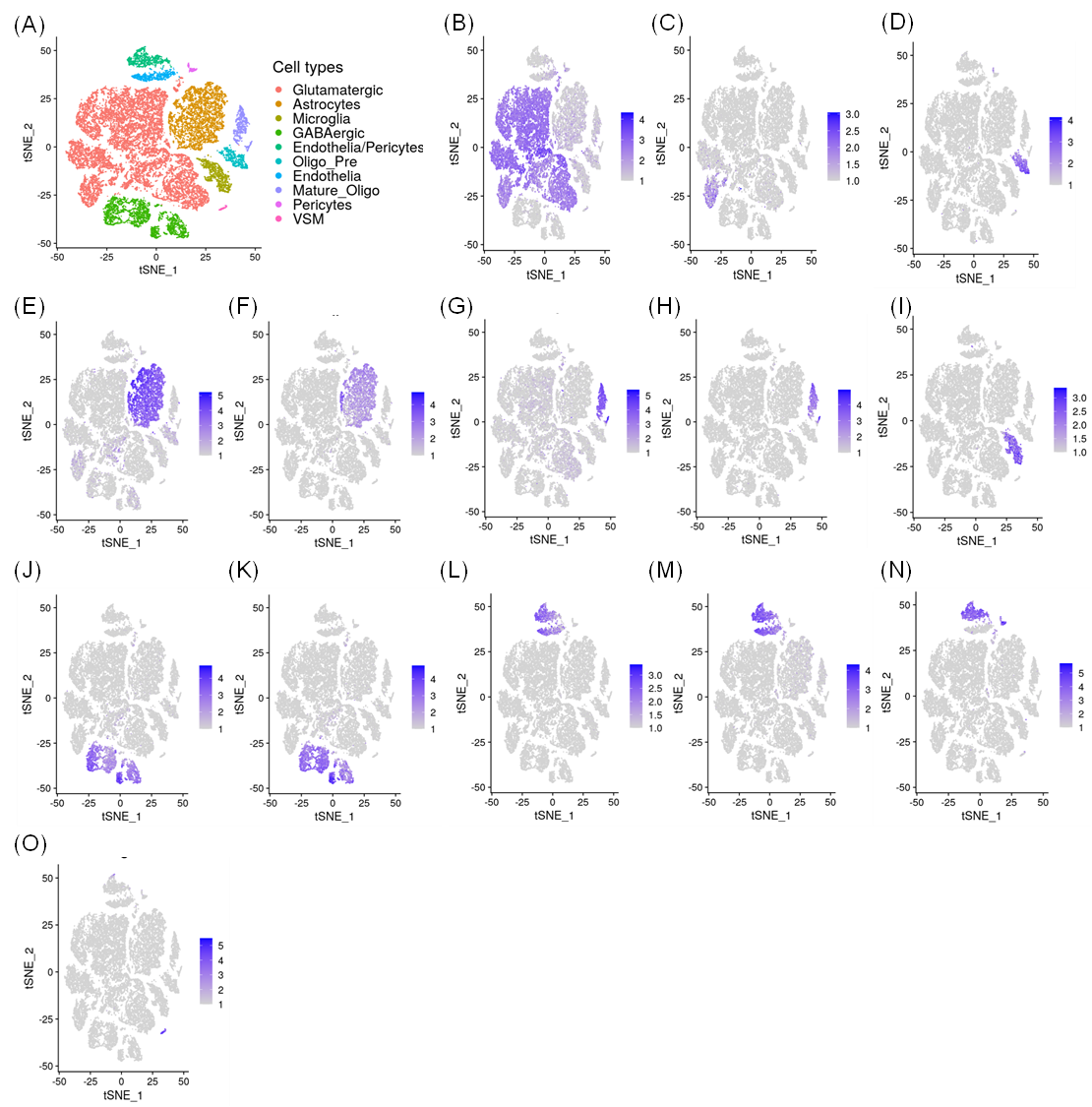

### Figure S5: Clustering by tSNE and cell type assignment

Data was clustered by tSNE and cluster identity was determined by normalized expression of cell specific markers. (A) Identity of clusters. Glutamatergic neurons determined by (B) *Slc17a7* and (C) *Slc17a6*. Oligodendrocytes precursors determined by (D) *Pdgfra*. Astrocytes determined by (E) *Aldoc* and (F) *Aqp4*. Mature oligodendrocytes determined by (G) *Mbp* and (H) *Cldn11*. Microglia identified by (I) *Tmem119*. GABAergic neurons identified by (J) *Gad1* and (K) *Gad2*. Endothelia determined by (L) *Pecam1* and (M) *Slc2a1*. Pericytes identified by (N) *Vtn*. Vascular smooth muscle identified by (O) *Tagln*.

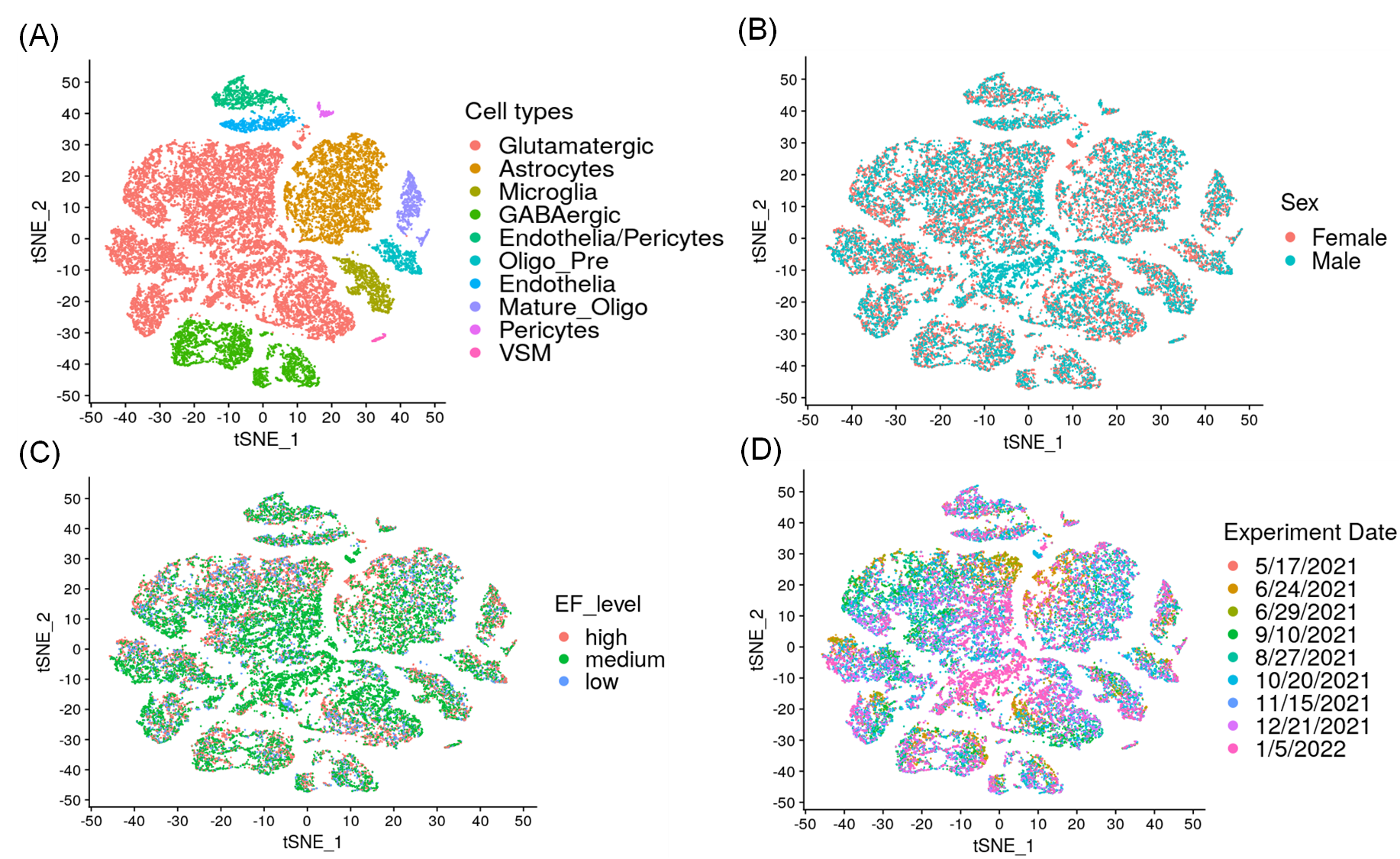

### Figure S6: Evaluating the effects of covariates on tSNE clustering

Cells were clustered by tSNE. (A) Clusters were identified by cell types. Overlay of covariates on tSNE by (B) Sex, (C) EF level and (D) Experiment date.

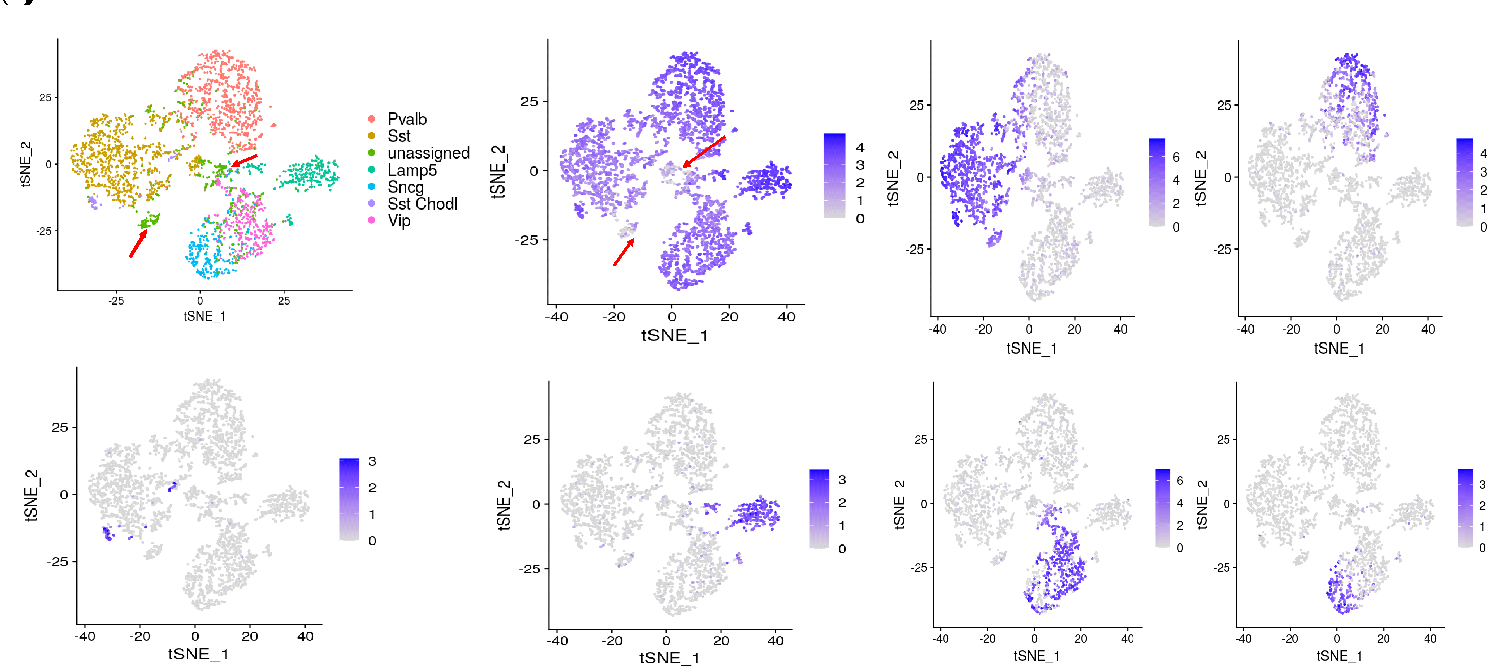
Figure S7: Cell type assignment of GABAergic neurons

GABAergic neurons were sub-clustered by tSNE and cell identities were determined using cell2cluster program with a reference PFC 10X dataset. (A) tSNE clustering of GABAergic neuron subtypes. Normalized expression of GABAergic cell type specific markers (B) *Gad1*, (C) *Sst*, (D) *Pvalb*, (E) *Chodl*, (F) *Lamp5*, (G) *Vip* and (H) *Sncg*. Red arrows highlight cells that had low or no gene expression of *Gad1* and were not assigned any cell identity by cell2cluster. These cells were therefore removed from further analysis.

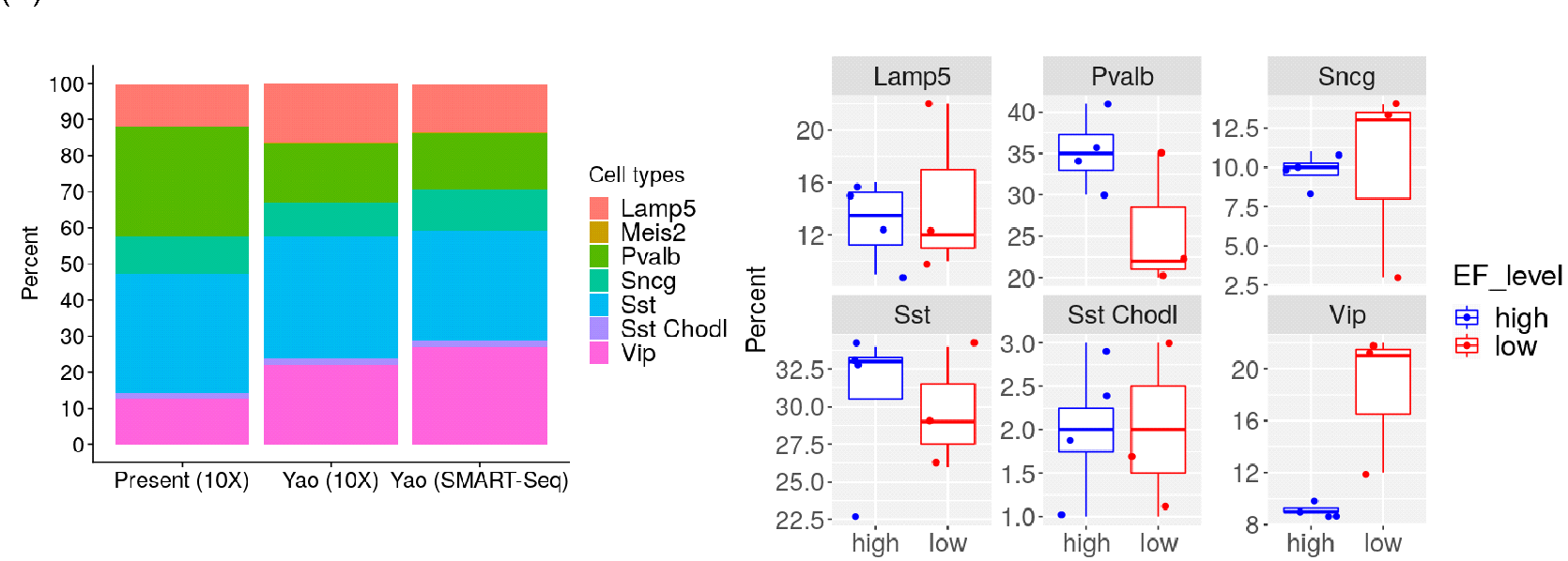

### Figure S8: Distribution of GABAergic neuron subtypes

A) Distribution of GABAergic neurons for this experiment and those from Yao et al, 2021 [2] processed by 10X and SMART-Seq. B) Comparison of GABAergic neurons between EF1 groups. Lamp5; p = 0.71, Pvalb; p = 0.17, Sncg; p = 0.95, Sst; p = 0.77, Sst Chodl; p = 1, Vip; p = 0.10.

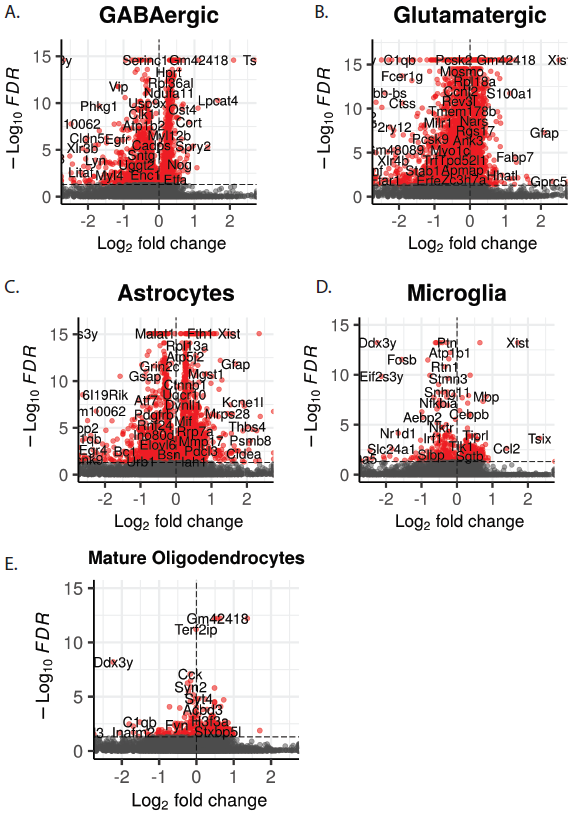

### Figure S9: Differential Gene Expression using DESingle pipeline

Volcano plots demonstrating significantly differentially expressed genes (padj<.05) in red for A) GABAergic inhibitory neurons B) glutamatergic neurons C) astrocytes D) microglia E) mature oligodendrocytes.

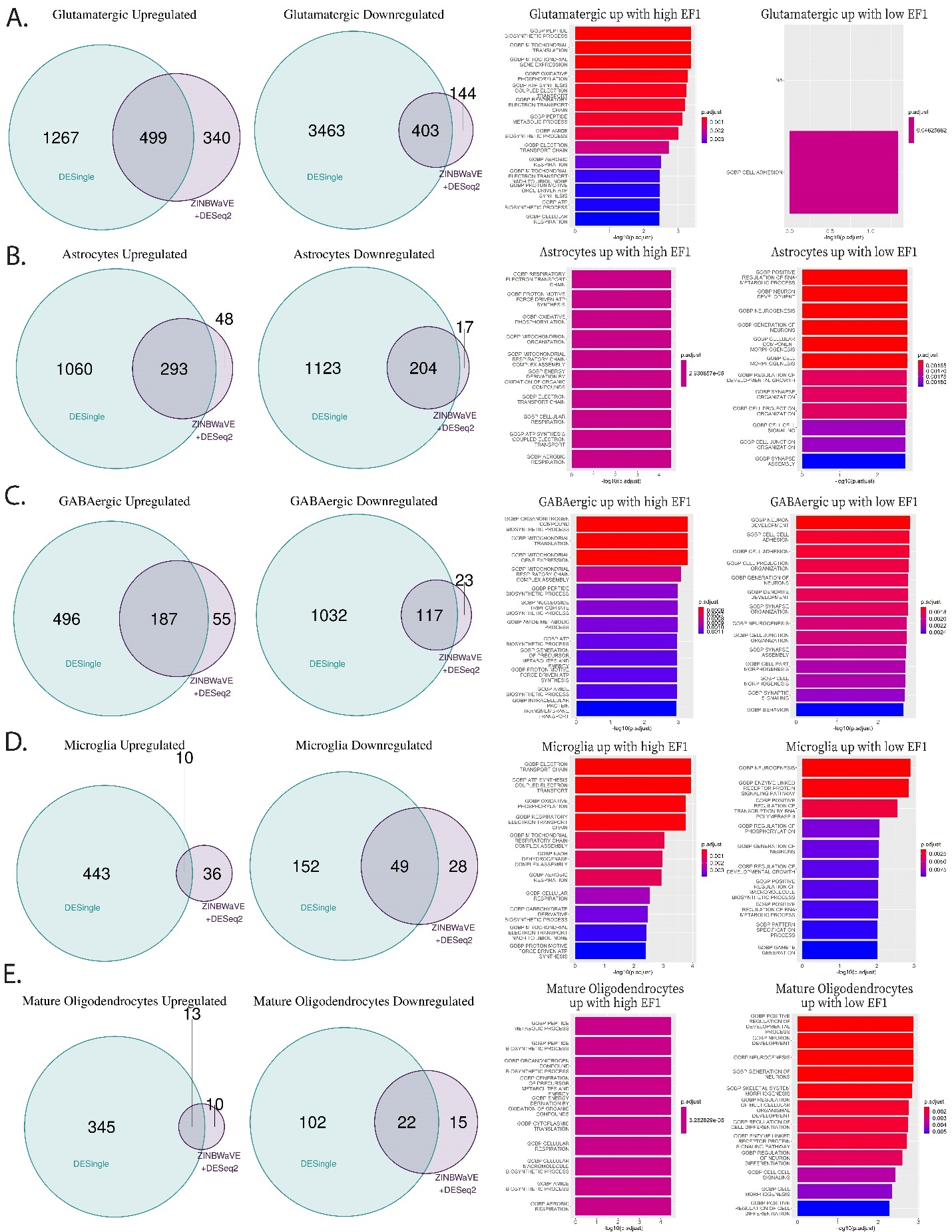

Figure S10: Validation of ZINB-Wave Findings using DESingle**.** For each cell type, from left to right is shown Venn Diagrams of overlapping genes between ZINB-Wave and DESingle, upregulated with EF1 (left), downregulated with EF1 (right), GSEA pathway analysis for enriched in genes with high EF1 (left), enrichment in genes for low EF1 (right). Data are show for A) glutamatergic neurons B) astrocytes C) GABAergic neurons D) microglia and E) mature oligodendrocytes.

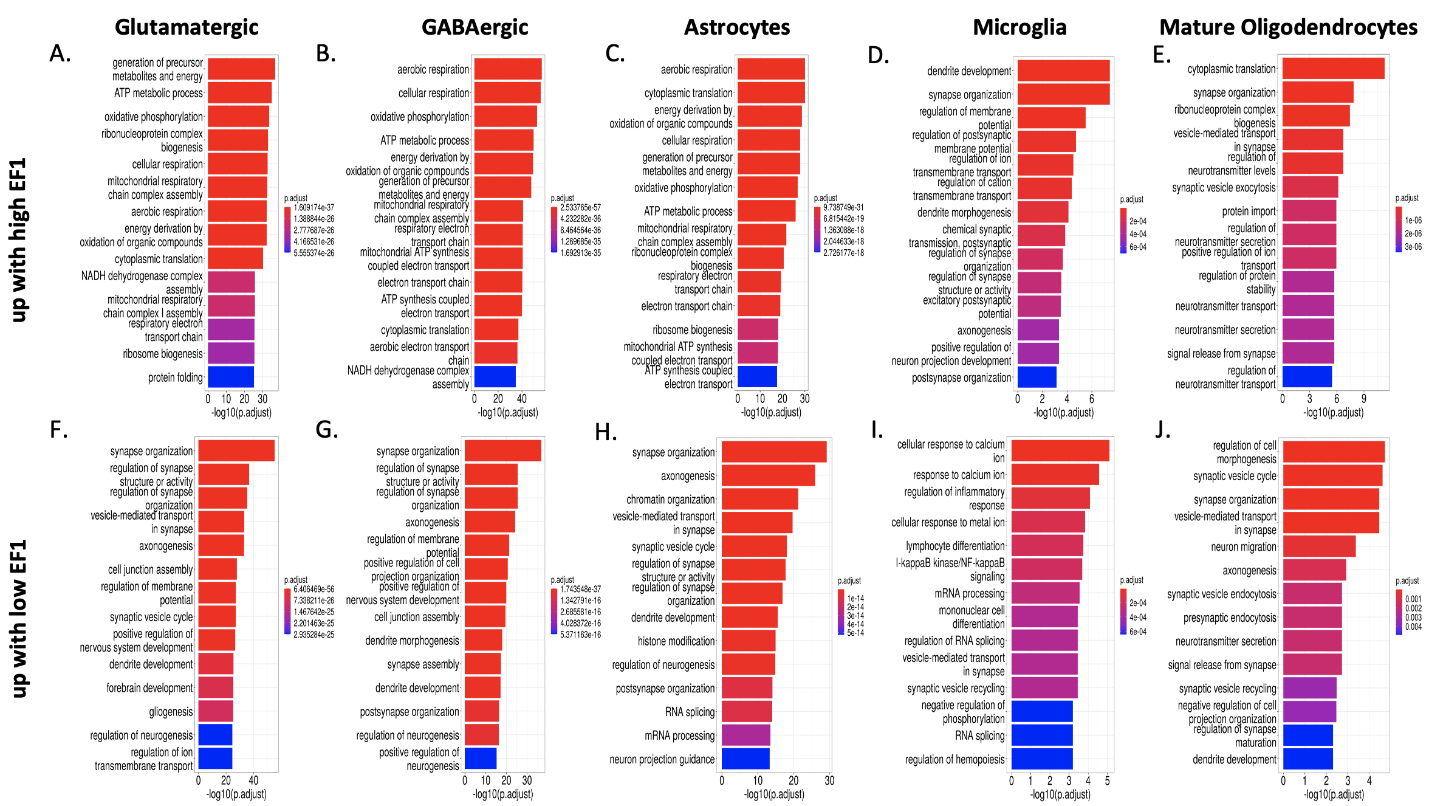

### Figure S11: GO Analysis using DESingle pipeline

For genes that were upregulated in mice that had high EF1 scores, GO terms were identified that were significantly enriched in (A) glutamatergic neurons, (B) GABAergic neurons, (C) astrocytes, (D) microglia and (E) mature oligodendrocytes. For genes that were upregulated in mice that had low EF1 scores, GO terms were identified that were significantly enriched in (F)glutamatergic neurons, (G) GABAergic neurons, (H) astrocytes, (I) microglia, and (J) mature oligodendrocytes.

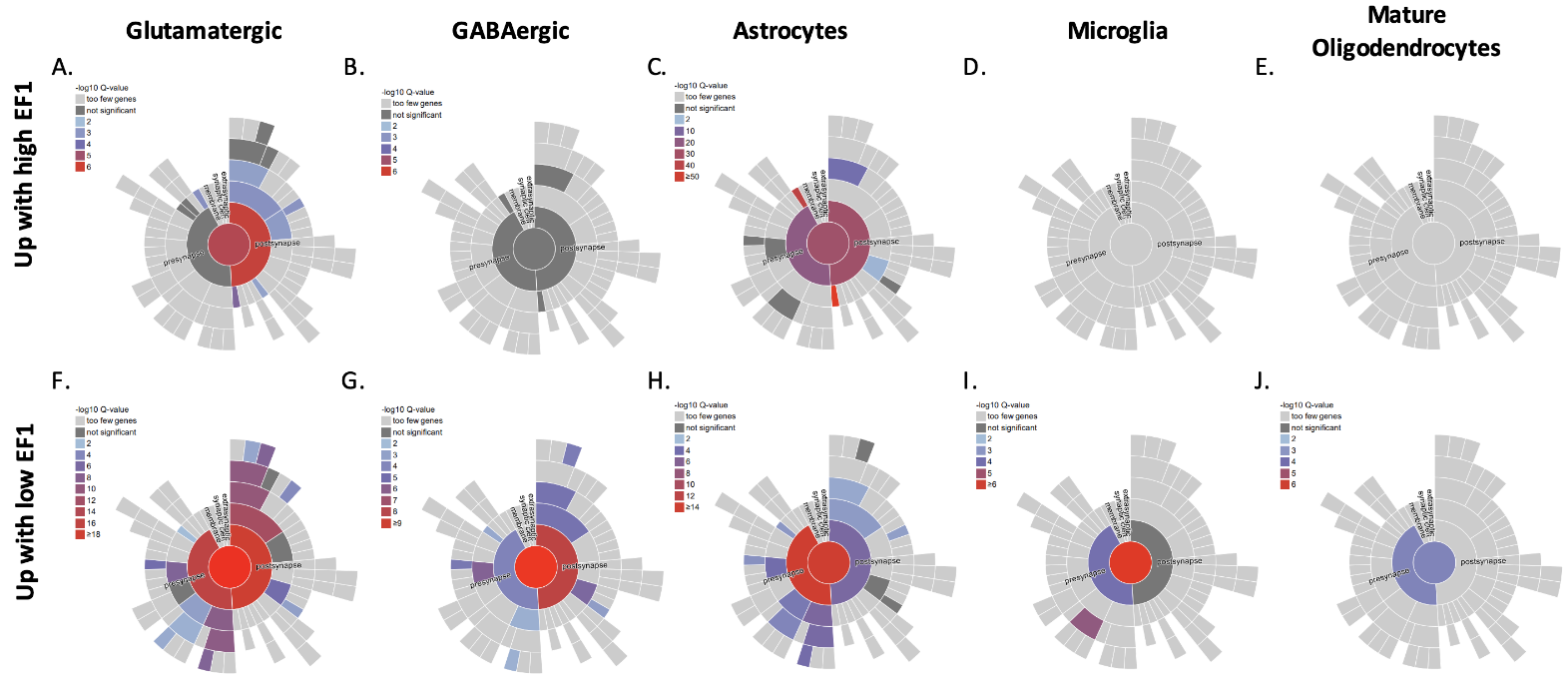

### Figure S12: SynGO over-representation analysis

For genes more highly expressed in the high EF1 group, over-representation for biological processes and cellular components related to synapse function was observed to a lesser degree for (A) glutamatergic neurons, (B) GABAergic neurons, and (C) astrocytes, and not observed at all for (D) microglia and (E) mature oligodendrocytes. For genes more highly expressed in the low EF1 group, significant over-representation for biological processes and cellular components related to synapse function was observed for all cell types (F-J).

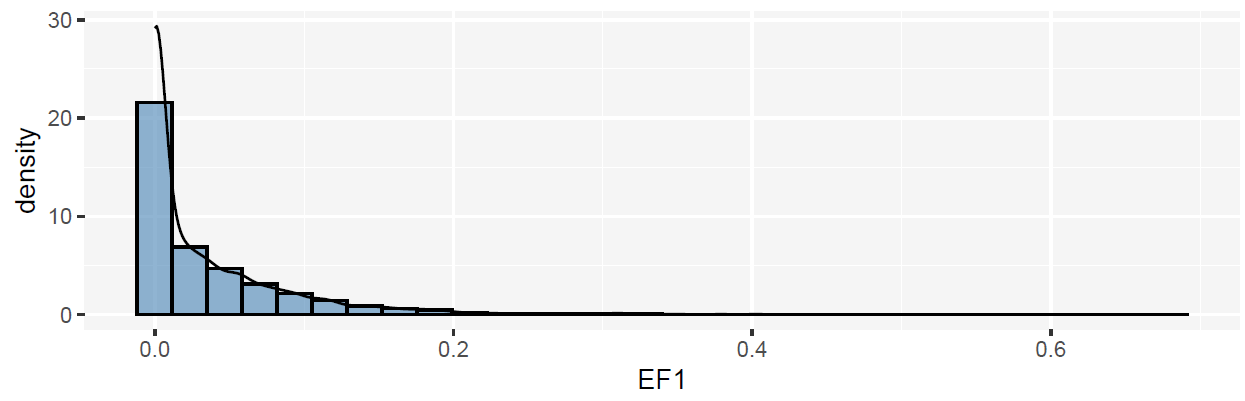

### Figure S13: Density estimate of the *Electome Factor* 1 scores on third portion of FIT

Density estimates of EF1 scores from 85 animals averaged across the 3^rd^ portion of the forced interaction test (FIT) (see also, Figure 1). Mean: 0.0362, Median: 0.0111, Standard Deviation: 0.0572.

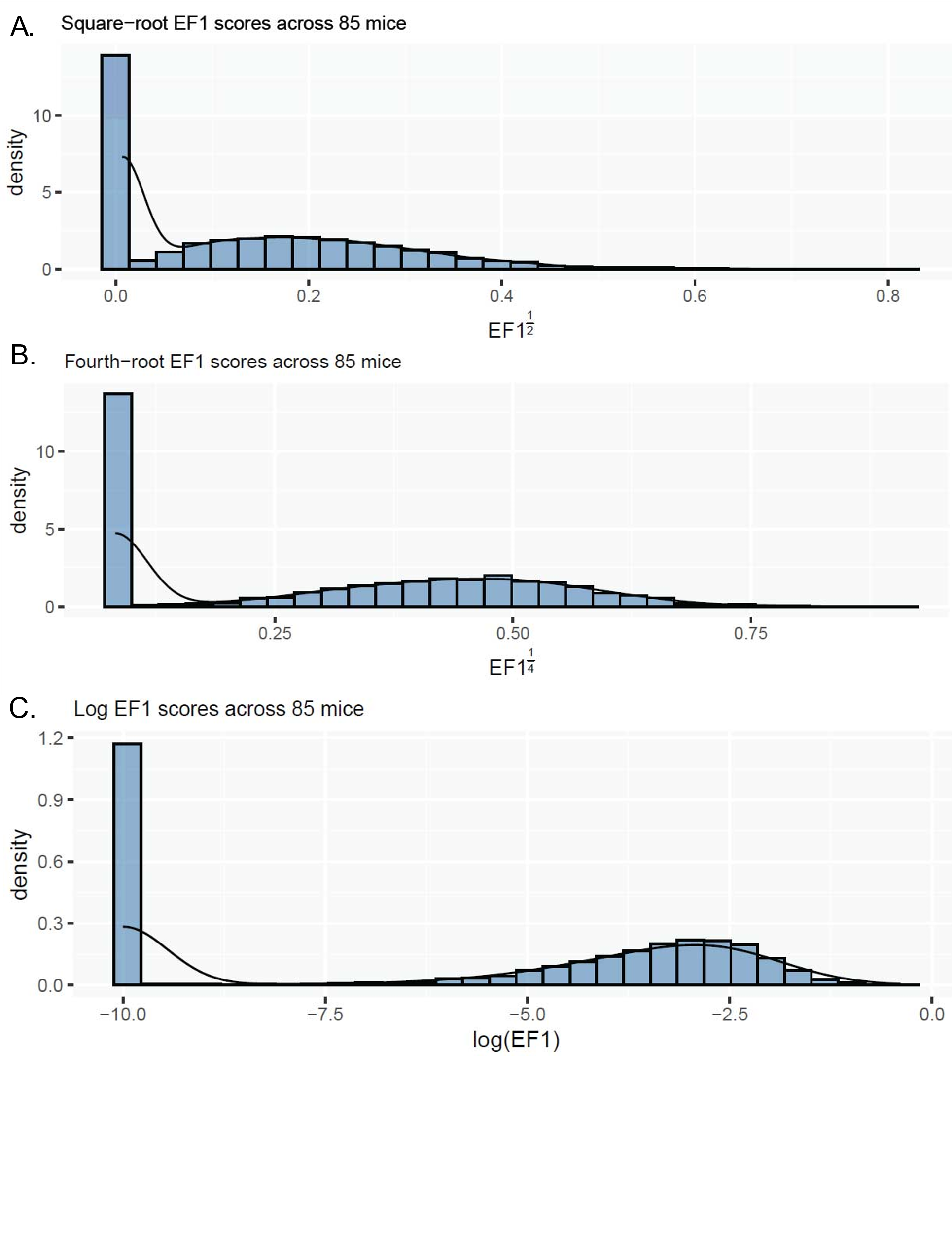

### Figure S14: Transformed densities of the *Electome Factor* 1 scores

Distributions of transformed *Electome Factor* 1 scores on the third portion of the FIT using A) square-root B) fourth-root and C) log transformations.

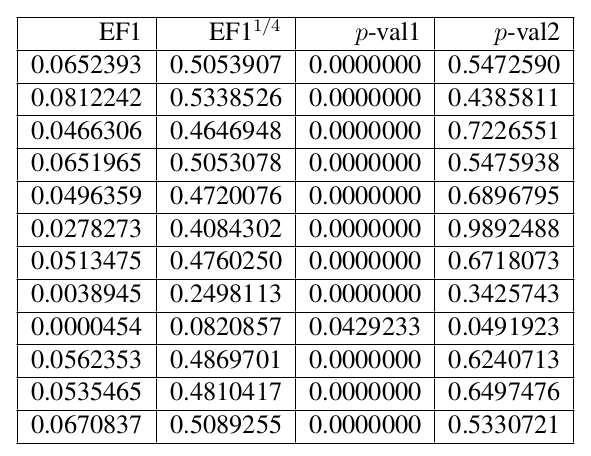

Figure S15: Mixed Normal p-values

This figure shows the p-values corresponding to the hypotheses that a given fourth-root transformed EF1 score comes from either the lower or upper part of the mixture of normal distribution discussed in the section “Electome Factor 1 Score Distributions”.

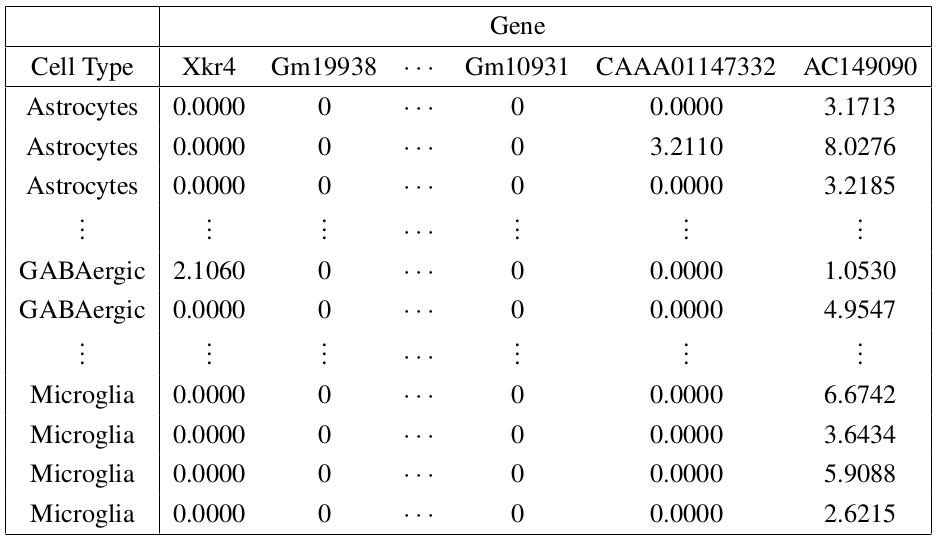

Figure S16: Example Full Cell Count Data

An example of a single mouse’s full cell count data post normalization. Note that the cell types have multiple replications for the same gene.

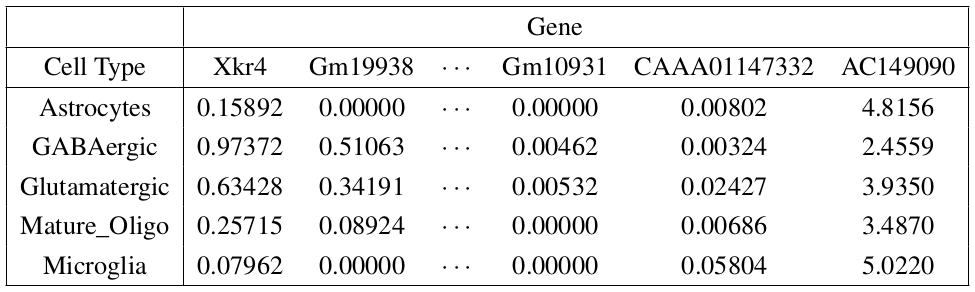

Figure S17: Example Mean Cell Count Matrix

An example of a single mouse’s mean cell count matrix, formed by taking the mean across all gene cell count vectors corresponding to each of five cell types. Note that each cell type and gene combination appears only once.

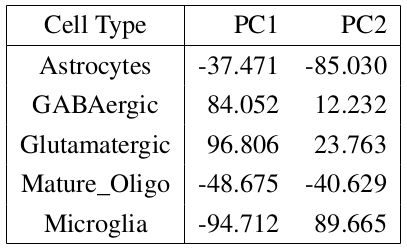

Figure S18: Example Principal Components Matrix

An example of a single mouse’s covariates, computed as the first two principal components of its mean cell count matrix, an example of which can be seen in Supplementary Figure S16.

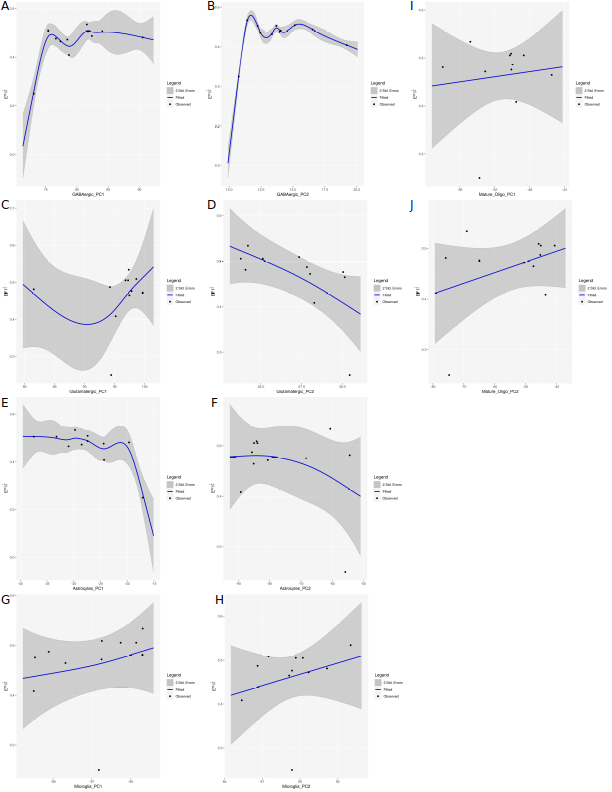

### Figure S19: Relationship between cell types and fourth root EF1

The GAM model estimates of the fourth-root *EF*1 score as a function of the first and second principal components for each cell type. The blue curve represents the estimate of the regression function and the gray bands denote the pointwise two standard deviation confidence intervals. The regression function and the uncertainty estimates are computed using the mgcv R package [4].

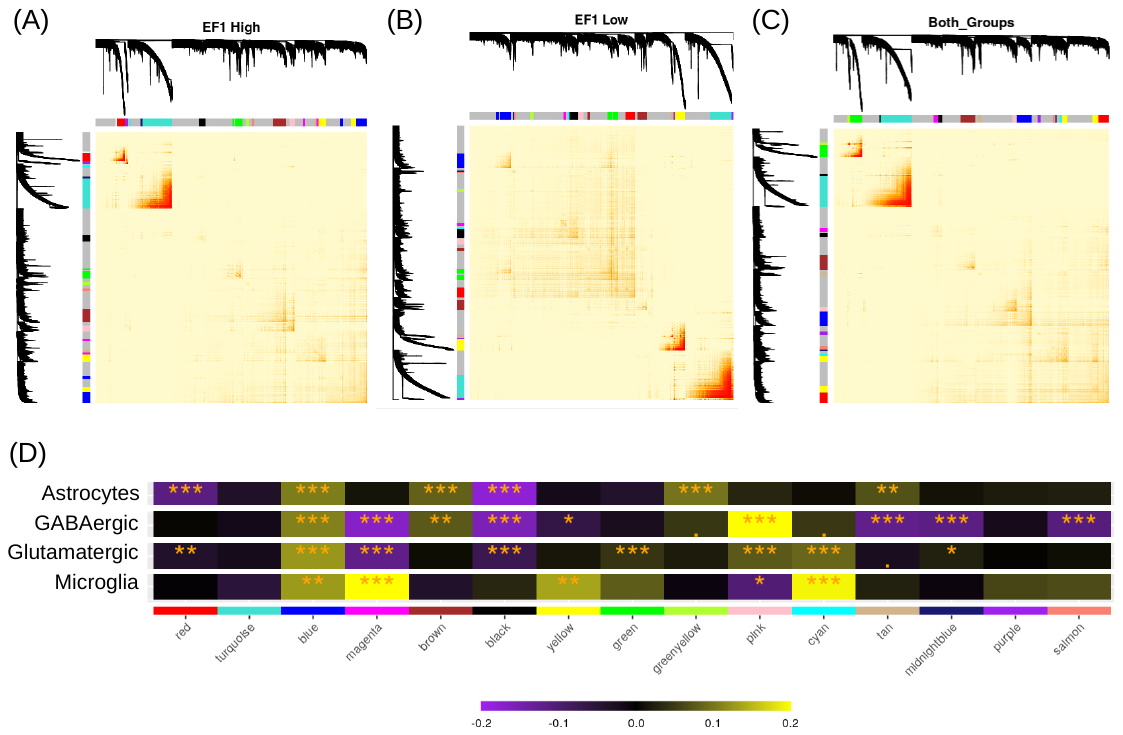

### Figure S20: Heatmap of Module-Trait correlation strength

Module-Traits correlation between eigengene of modules with *EF*1 groups. *; p < 0.05, **; p < 0.01, ***; p < 0.001. Yellow indicates a positive correlation between eigengene and *EF*1 activity; purple indicates a negative correlation.

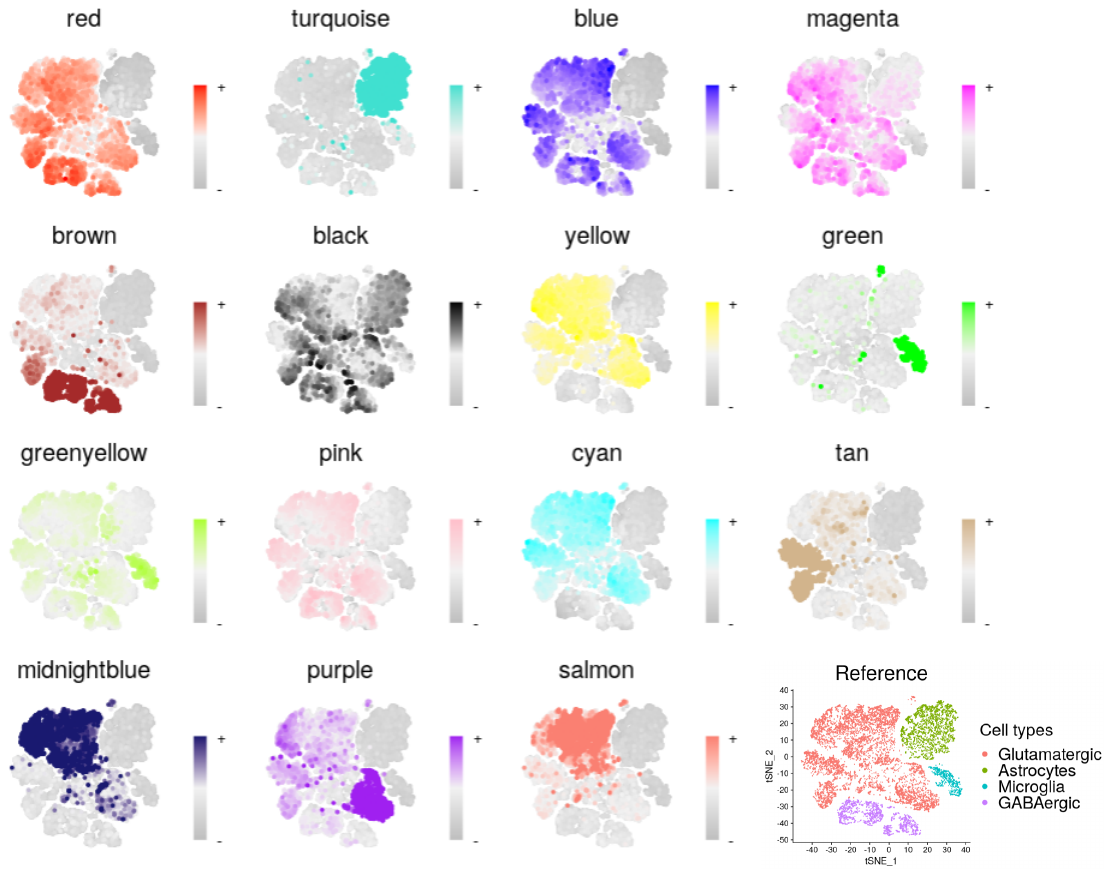

### Figure S21: Expression of module eigengenes across different tSNE clusters.

Eigengene expression for each module was overlayed on tSNE clusters. Positive eigengene expression are in the respective colors while negative eigengene expression are in dark grey.

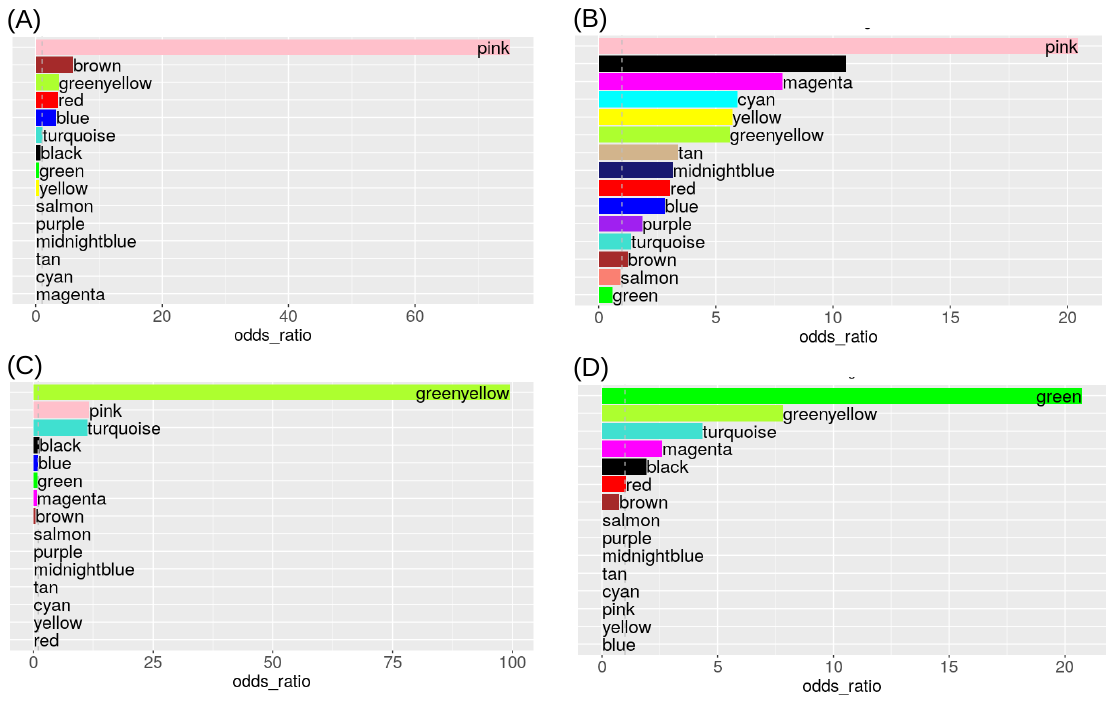
Figure S22: Enrichment of differentially expressed genes in modules. Modules were assessed for enrichment of differentially expressed genes from (A) GABAergic neurons, (B) glutamatergic neurons, (C) astrocytes and (D) microglia. Modules above the dotted line for each panel were enriched for differentially expressed genes that were more highly expressed in the high *EF*1 group. Genes that were reduced in the high *EF*1 group did not show enrichment for any of the modules.

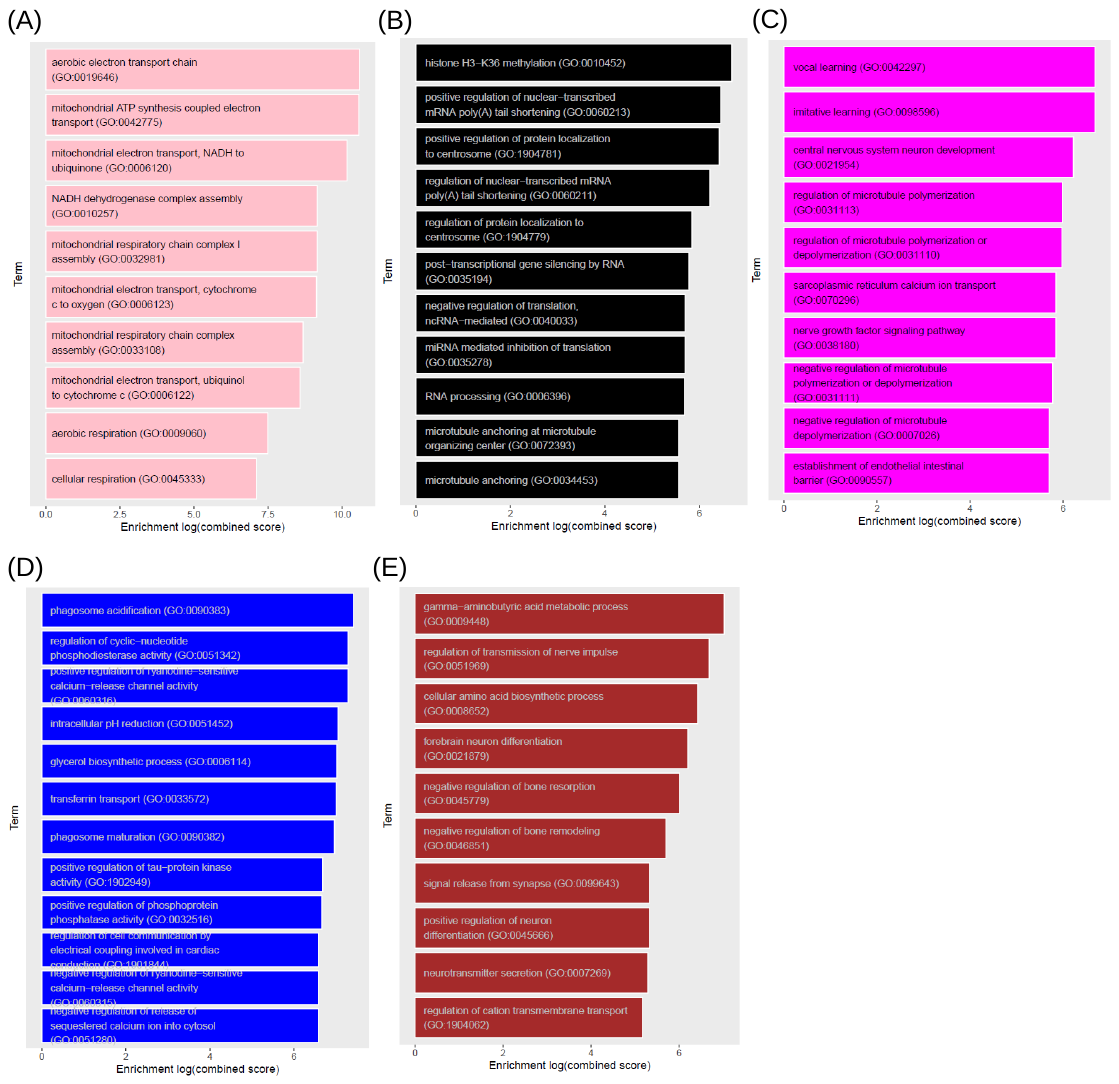

Figure S23: Module Gene Ontology analysis. Genes from each module that were significantly associated with *EF*1 group were used for gene ontology analysis to identify biological processes significantly over-represented by these genes. Terms showed are the top 10 terms arranged by combined score that survived multiple correction. Modules shown are (A) pink, (B) black, (C) magenta, (D) blue, and (E) brown.

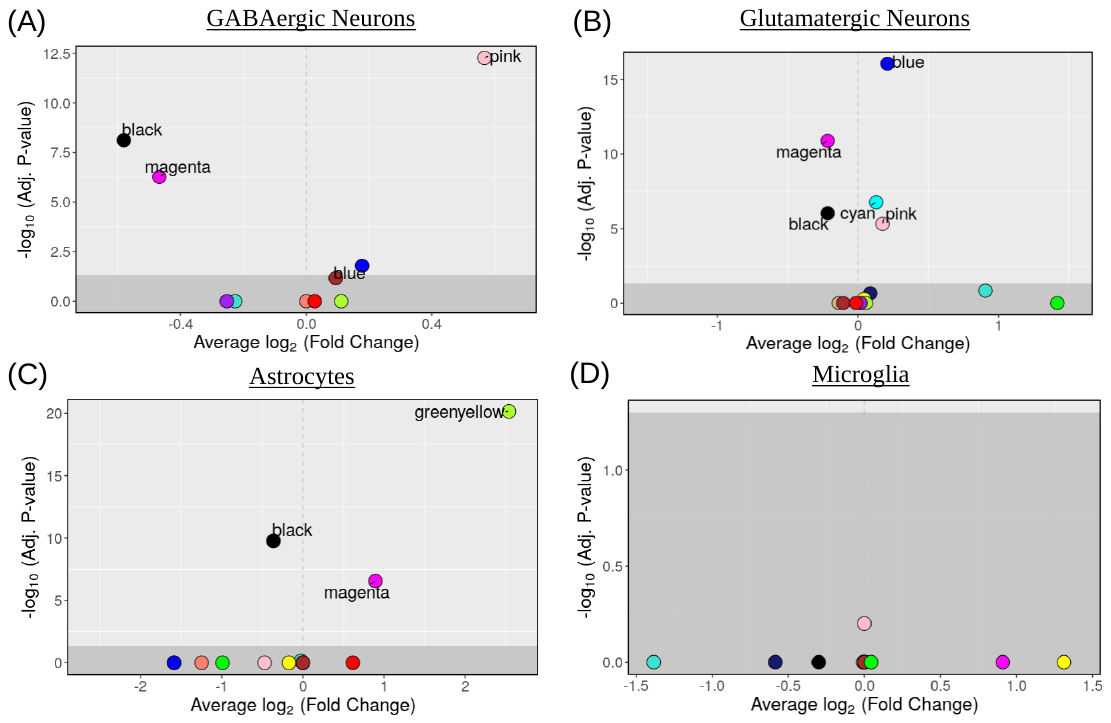

Figure S24: Volcano plot of differential eigengene expression.

Eigengene expression for each module was compared between high and low *EF*1 groups for (A) GABAergic neurons, (B) Glutamatergic neurons, (C) Astrocytes and (D) Microglia. Modules in the light grey shaded area are modules that survived Bonferroni multiple correction (Adjusted P-value < 0.05). Fold change > 1 indicates higher expression in high *EF*1 group compared to the low *EF*1 group. Fold change < 1 indicates lower expression in the high *EF*1 group compared to the low *EF*1 group.

### Table S1: Covariate Information

See Table attachment “Supplemental_Table_S1_Covariate_Information.xlsx”

### Table S2: Cell Ranger Alignment Statistics

See Table attachment “Supplemental_Table_S2_Cell_Ranger_Aligement_Statistics.csv”

### Table S3: Table of Cell Types and Numbers

See Table attachment “Supplemental_Table_S3_Final_Cell_Count.csv”

### Table S4: Table of Cell Type Proportions

See Table attachment “Supplemental_Table_S4_Final_Cell_Proportions.csv”

### Table S5: GSEA and GO Biological Processes

See Table attachment “Supplemental_Table_S5_GSEA_GO_BP.xlsx”

### Table S6: Mitochondrial GSEA

See Table attachment “Supplemental_Table_S6_GSEA_mito.xlsx”

### Table S7: DEGs for ZINB-WavE analysis with annotations

See Table attachment “Supplemental_Table_S7_ZINBWavE_analysis_annotations.xlsx”

### Table S8: ZINB-WaVE Differentially Expressed Genes in Major Cell Types

| **Number of genes with padj < .05** | | |
| --- | --- | --- |
| **Cell Type** | **up with high *EF1*** | **up with low *EF1*** |
| Glutamatergic Neurons | 838 | 548 |
| Astrocytes | 341 | 221 |
| GABAergic Neurons | 242 | 140 |
| Microglia | 46 | 77 |
| Mature Oligodendrocytes | 23 | 37 |

### Table S9: ZINB-WaVE Differentially Expressed Genes in GABAergic Neurons

| **Number of genes with padj < .05** | | |
| --- | --- | --- |
| **Cell Type** | **up with High *EF1*** | **up with Low *EF1*** |
| Sst | 73 | 129 |
| Pvalb | 2 | 4 |

### Table S10: DESingle Differential Gene Expression in Major Cell Types

| **Number of genes with padj < .05** | | |
| --- | --- | --- |
| **Cell Type** | **up with High *EF1*** | **up with Low *EF1*** |
| Glutamatergic Neurons | 1824 | 3808 |
| Astrocytes | 1304 | 1376 |
| GABAergic Neurons | 687 | 1145 |
| Microglia | 441 | 213 |
| Mature Oligodendrocytes | 327 | 155 |

### Table S11: DESingle Differential Gene Expression in GABAergic subtypes

| **Number of genes with padj < .05** | | |
| --- | --- | --- |
| **Cell Type** | **up with High *EF1*** | **up with Low *EF1*** |
| Sst | 426 | 315 |
| Pvalb | 127 | 39 |

### Table S12: DESingle Main Cell Types DEG

See Table attachment “Supplemental_Table_S12_DESingle_Main_Cell_Types_DEG.xlsx”

### Table S13: DESingle Main Cell Types GO

See Table attachment “Supplementary_Table_S13_DESingle_Main_Cell_Types_GO.xlsx”

### Table S14: DESingle ZINB-WaVE Overlap

See Table attachment “Supplementary_Table_S14_DESingle_ZINB-WaVE_Overlap.xlsx”

### Table S15: Upstream Regulator Analysis

See Table attachment “Supplementary_Table_S15_Upstream_Transcriptional_Regulators.xlsx”

### Table S16: Gaussian Process Model Performance

| **model_id** | **n_pc** | **Degree** | **train_err** | **test_err_mean** | **test_err_sd** |
| --- | --- | --- | --- | --- | --- |
| 1 | 1 | 1 | 0.308732402 | 0.009182841 | 0.00165257 |
| 2 | 1 | 2 | 0.032570205 | 0.01047314 | 0.002813304 |
| 3 | 2 | 1 | 0.213841216 | 0.010470759 | 0.001839324 |
| 4 | 2 | 2 | 0.001600662 | 0.011444574 | 0.001369333 |

### Table S17: Min-Max *Electome Factor* 1 (4^th^ root) in Gaussian Process Model

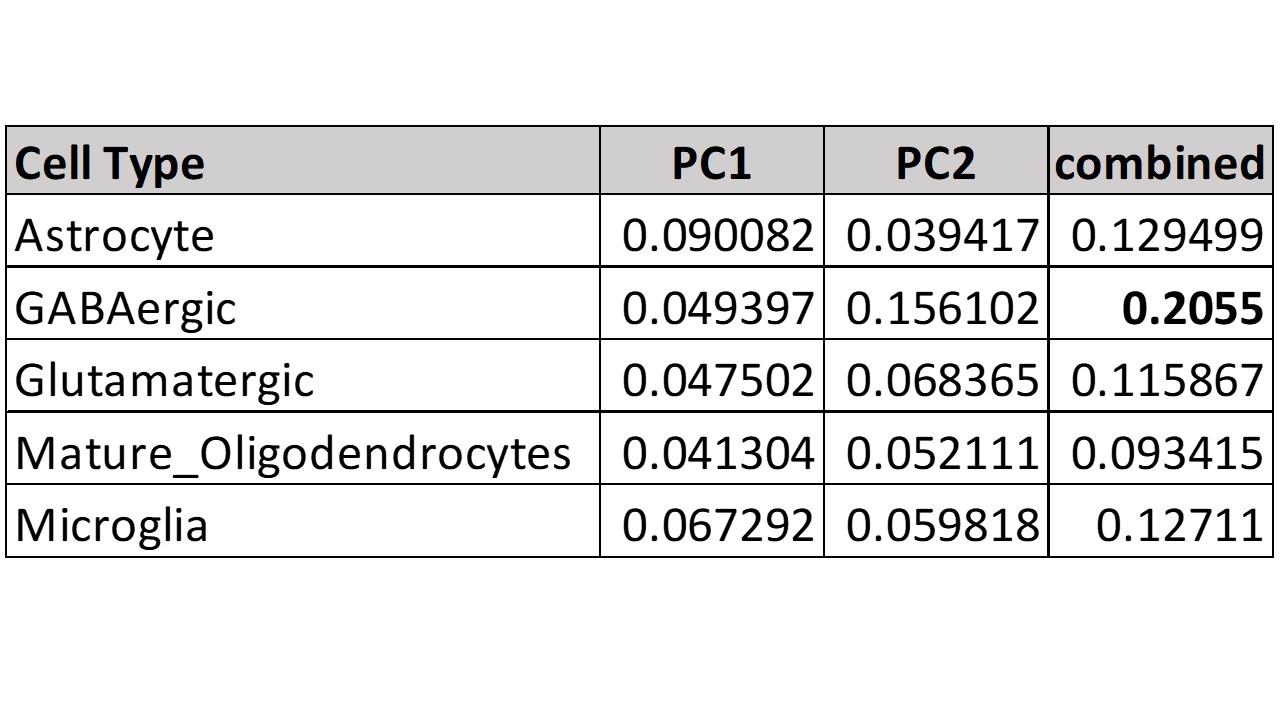

### Table S18: Comparison with Chronic Social Defeat Stress Data (Susceptibility)

See Table attachment “Supplementary_Table_S18_Bagot Susceptibility Module overlap with DEGs by cell type.xlsx”

### Table S19: Comparison with Chronic Social Defeat Stress Data (Resilience)

See Table attachment “Supplementary_Table_S19_Bagot Resilience Module overlap with DEGs by cell type.xlsx”

### Table S20: Comparison with Hub Genes from Chronic Social Defeat Stress Data

See Table attachment “Supplementary_Table_S20_DEG_overlap_with_Bagot_midnightblue_and_violet_hub_genes.xlsx”

### Table S21: MAGMA

See Table attachment “Supplementary_Table_S21_MAGMA.xlsx”

### Table S22 Gene ontology analysis of modules

See Table attachment “Supplementary_Table_S22_Gene_Ontology_Analysis_Modules.xlsx”

### Table S23: Differential connectivity of reactome pathways

See Table attachment “Supplementary_Table_S23_Differential_Connectivity_Reactome_Pathway.xlsx”

Mixture Models.” Journal of Statistical Software, **32**(6), 1–29.

<https://www.jstatsoft.org/v32/i06/>.
